# Heterologous expression of lyngbyatoxin biosynthetic genes in *Aspergillus oryzae* reveals transcriptional barriers but enables LtxC-mediated biotransformation

**DOI:** 10.64898/2026.05.15.725566

**Authors:** Sameera Jayasundara, Tahir Ali, Bisola Adeyemi, Bagyashri Krishnamorthy, Calvin A. Henard, Kent D. Chapman, Elizabeth Skellam

## Abstract

Cyanobacterial natural products are a rich source of bioactive compounds, yet their heterologous production remains challenging. This study investigates the feasibility of expressing the lyngbyatoxin A (LTXA) biosynthetic gene cluster in a fungal host. The lyngbyatoxin biosynthetic genes (*ltxA, ltxB, ltxC*) were individually cloned and expressed in *Aspergillus oryzae* NSAR1 under the control of an inducible promoter. Metabolite production was assessed using LC– MS, and transcriptional analysis was performed by RT-PCR. Codon-optimized constructs and precursor feeding experiments were employed to evaluate pathway functionality. No production of LTXA or pathway intermediates was detected upon co-expression of *ltxA–C* despite confirmed transcription of *ltxB* and *ltxC*. RT-PCR analysis revealed truncation of the *ltxA* transcript, suggesting incompatibility with fungal transcriptional or splicing machinery. In contrast, expression of a codon-optimized *ltxC* enabled biotransformation of indolactam V to LTXA in *A. oryzae*, confirming functional expression of the prenyltransferase. These results highlight transcriptional limitations as a key barrier to heterologous expression of cyanobacterial NRPS pathways in fungal hosts, while demonstrating that downstream tailoring enzymes can remain functional. This work provides insights for future engineering of fungal platforms for cyanobacterial natural product biosynthesis.

## Background

Cyanobacterial natural products represent a diverse source of bioactive molecules with pharmaceutical and biotechnological potential (Welker & Von Dohren, 2006; Singh et al., 2011; Calteau et al., 2014) (Figure 1). However, access to these compounds is often limited by the slow growth of native producers and the lack of robust genetic tools (Philmus et al., 2026), making heterologous expression an attractive strategy for their production and study (Berla et al., 2013; Baunach et al., 2023). Despite advances in pathway engineering, heterologous expression of cyanobacterial biosynthetic gene clusters remains challenging, (Cullen et al., 2019; Dhakal et al., 2021) particularly for large multi-domain non-ribosomal peptide synthetase (NRPS) systems. Differences in codon usage, transcriptional regulation, and protein folding between hosts can hinder functional expression, and successful reconstitution of complete pathways is rare (Ongley et al., 2013; Videau et al., 2020; Chen et al., 2025; Philmus et al., 2026).

**Figure 1.**
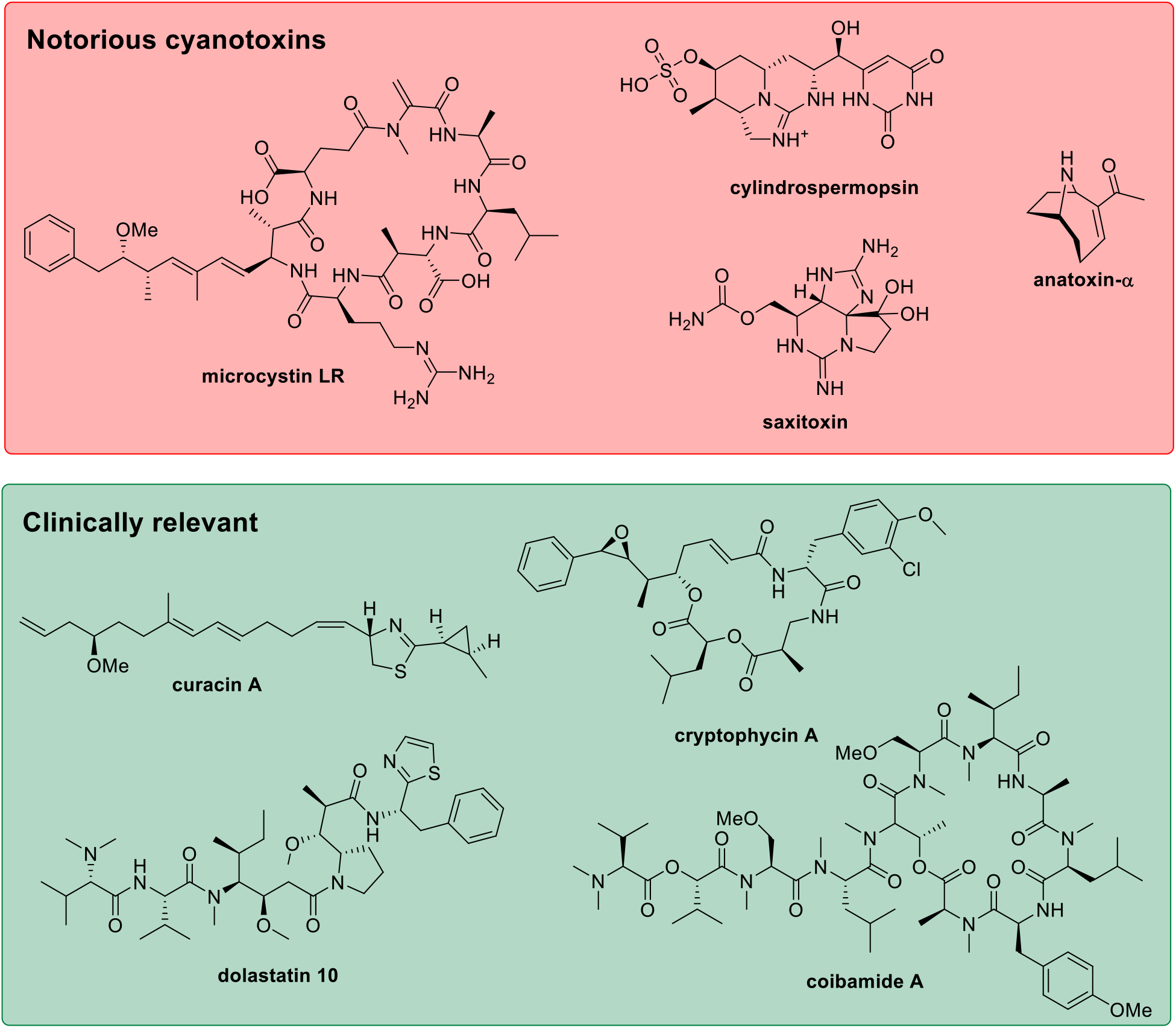
Examples of infamous and clinically relevant natural products of cyanobacterial origin.

Fungal hosts such as *Aspergillus oryzae* are attractive heterologous expression host alternatives due to their genetic tractability and established use in natural product biosynthesis (Lazarus et al., 2014; Alberti et al., 2017; Feng et al., 2021) but their suitability for expressing cyanobacterial pathways remains underexplored. In this study, we investigated the heterologous expression of the well-characterized lyngbyatoxin A (LTXA) biosynthetic gene cluster (BGC; *ltxA-C;* Scheme 1) in *A. oryzae*. LTXA is a protein kinase C (PKC) activator requiring valine, tryptophan, and geranyl pyrophosphate (GPP) precursors (Edwards & Gerwick, 2004) which are readily available in *A. oryzae* NSAR1; the GC content of the *ltx* genes (∼45%) is similar to that of *Aspergillus* sp. genes (∼52%); and the expression system enables individual cloning of each gene under the control of a strong fungal inducible promoter (Lazarus et al., 2014). Our results reveal transcriptional limitations affecting NRPS expression, while demonstrating that downstream tailoring enzymes can remain functional in a fungal system.

**Scheme 1:**
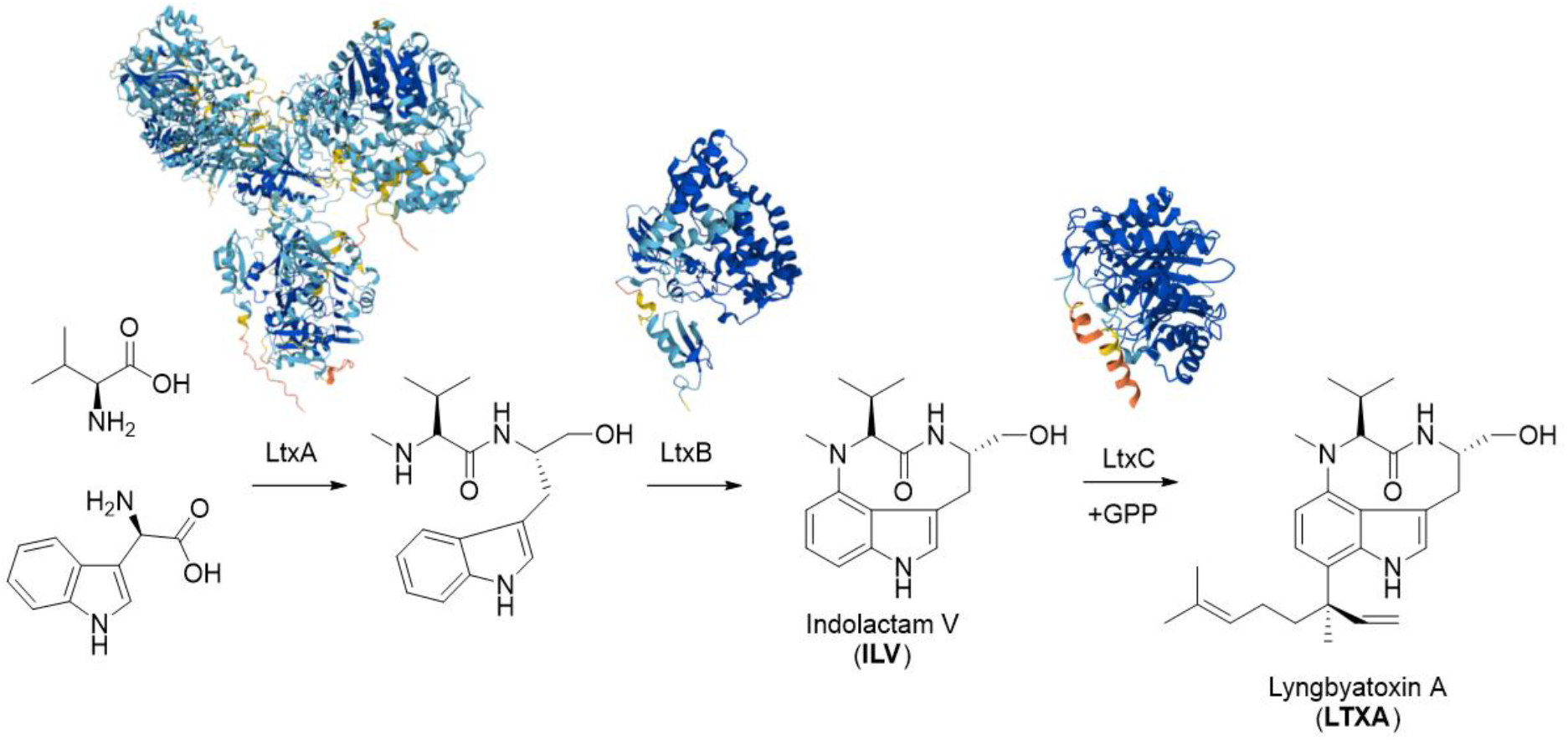
Biosynthesis of LTXA. The NRPS gene *ltx*A encodes the following domain organization A-NM-T-C-A-T-R, where A = adenylation, NM = N-methyltransferase, T = thiolation, C = condensation, R = reductase.

## Results

### Cloning and heterologous expression of the *ltx* BGC

As PKC is a crucial signaling enzyme in fungi and potential target for anti-fungal drugs (Schmitz & Heinisch, 2003; Lafayette et al., 2010), the sensitivity of wild-type *A. oryzae* NSAR1 was established by cultivating in the presence of up to 200 μg LTXA. Over the seven day observation period, *A. oryzae* NSAR1 grew normally and organic extracts of the mycelium and culture broth confirmed that LTXA was not modified by native enzymes. Based on three replicate extractions, over 90% of the added LTXA sample could be recovered from cell samples, whereas less than 5% could be recovered from the culture broth (Table S10).

The three essential *ltx* genes were commercially synthesized and individually cloned under the inducible *amyB* promoter in the pTYGS series of expression plasmids (Pahirulzaman et al., 2012).. The three resulting plasmids were confirmed by sequencing and co-transformed using PEG-mediated protoplast transformation with a fourth plasmid containing a phosphopanthethienyl transferase (PPtase) - either the universal *sfp* PPtase from *Bacillus subtilis* or the *hetI* PPtase from *Nostoc* sp. PCC 7120 (Videau et al., 2016). Resulting transformants were selected three times on Czapek-Dox medium lacking arginine, adenine, methionine, and ammonium sulfate, before being transferred to liquid DPY medium. After seven days, cells were filtered from the culture broth and both were separately extracted twice using EtOAc and concentrated prior to LCMS analysis.

Comparing resultant extracts from transformants to commercial standards of LTXA and ILV using LCMS analysis revealed that none were producing the expected metabolites, known analogues (Soeriyadi et al., 2022), nor was there any indication of modification of these metabolites. RT-PCR on mRNA collected from transformants cultivated for 3 – 4 days confirmed that *ltxB* and *ltxC* were being transcribed, however, *ltxA* transcripts were determined to be truncated (Figure S6). Since the *amyB* promoter is inducible by starch or starch-derived oligosaccharides, and repressed by glucose, we adjusted our cultivation parameters in an attempt to modify transcription rate and timing (Table 1).

**Table 1:** Summary of fermentation optimization attempts. DPY = dextrin peptone yeast extract medium; CZDB = Czapek-Dox broth (+/-supplementation with 2% glucose or 2% starch); co = codon optimized.

| Strain | Conditions tested | Target metabolites detected | Outcome |
| --- | --- | --- | --- |
| *A. oryzae* (*ltxABC*+*sfp/hetI*) | Multiple media (DPY, CZDB ± glucose/starch), 3–12 days | None | *ltxA* transcript truncated; no precursors available |
| *A. oryzae* (*ltxABC*+*hetI*) + ILV | DPY, 7 days | None | No biotransformation despite all genes present and transcribed |
| *A. oryzae* (*ltxC-co*) + ILV | DPY, 7 days | LTXA ($m/z$ 438.3 = [M+H]<sup>+</sup>) | Successful biotransformation |

*In silico* analysis using GenScript’s rare codon analysis tool indicated that *ltxA* possesses negative *cis* elements (Jain et al., 2023; Fan et al., 2024), even when codon optimized for *Aspergillus* sp. Additionally, FGENESH analysis (Solovyev et al., 2006) indicated that *Aspergillus* sp. splice sites are detected in the *ltxA* native sequence (Figure S7) and challenges should be expected even if a codon optimized sequence is used (Figure S8). Therefore, we focused on understanding how to optimize production of LtxB and LtxC in a fungal host.

### Biotransformation of indolactam V by LtxC in *Aspergillus oryzae*

Since cyanobacterial gene sequences are known to utilize rare codons (Jones et al., 2012; Chen et al., 2025), *ltxC* was re-synthesized codon optimized (*ltxC*-co) for *Aspergillus* sp. and transformed into *A. oryzae* NSAR1. In parallel experiments, 100 μg of ILV was added to an individual *A. oryzae* + *ltxABC*/*ltxC*-co transformants in liquid DPY medium. After seven days the cultures were extracted and analyzed by LCMS. While ILV (*m/z* 302.1863 *=* [M+H]^+^) could be detected in the *A. oryzae* + *ltxC*-co media extract (Figure 2A), less was detected in the cell extracts (Figure 2B). Although LTXA (*m/z* 438.3115 = [M+H]^+^) was not detected in the media extracts of *A. oryzae* + *ltxC*-co (Figure 2C) it could clearly be detected in the cell extracts (Figure 2D). The production of LTXA in *A. oryzae* + *ltxC*-co transformants was confirmed by MS/MS analysis with a commercial standard (Figure 2E). In contrast, no conversion of ILV to LTXA was detected in *A. oryzae + ltxABC + hetI* strains (Figure S27).

**Figure 2:**
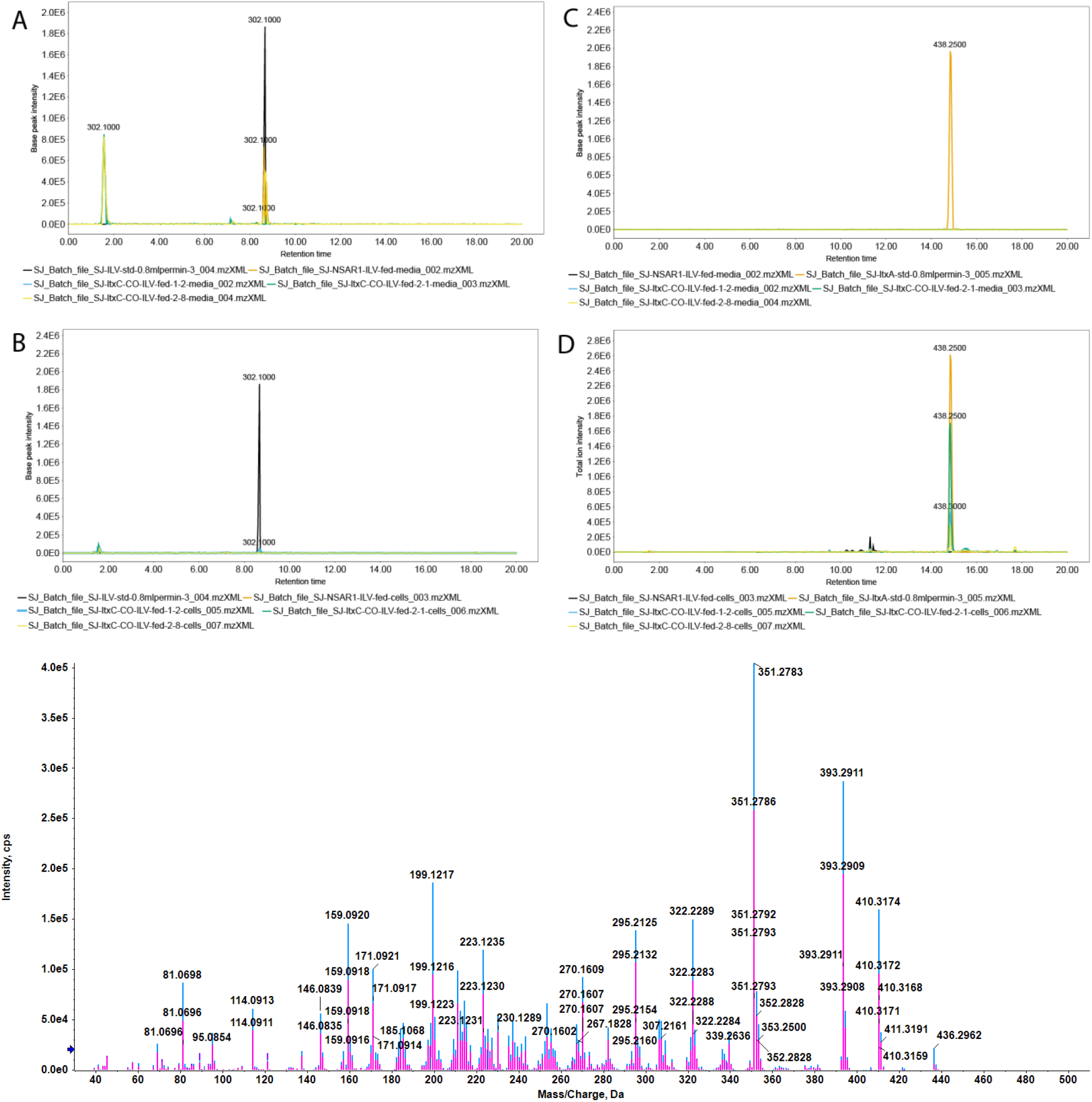
LCMS analysis of *A. oryzae* + *ltxC*-co transformant extracts after supplementation of ILV: A) media extract showing trace amounts of ILV; B) cell extracts showing no ILV; C) media extracts showing no LTXA; D) cell extracts showing LTXA production; E) MS/MS overlay of LTXA standard and the organic extract from a transformant. Lyngbyatoxin A standard (blue) and the cell extract of ILV-fed ltxC-Co transformants 21 (pink)

### Recombinant expression of LtxB does not indicate translational coupling

In the homologous *tle* BGC, which produces LTXA as an intermediate during teleocidin B in *Streptomyces blastmyceticus* (Zhang et al., 2016), the P450 enzyme and MbtH protein are encoded by two separate genes (Figure 3A-C). To investigate whether translational features of *ltxB* could contribute to pathway failure in a fungal host, LtxB was heterologously expressed in *Escherichia coli* with either N- or C-terminal 6xHis tags. SDS–PAGE analysis revealed a single protein band at ∼53 kDa for both constructs, and Western blot analysis confirmed expression of the C-terminally tagged protein. Peptide sequencing of purified LtxB confirmed that the correct protein had been recombinantly produced, however the N-terminal His-tagged protein may have lost its His-tag via Ni-NTA affinity column purification. No evidence for a smaller (∼9 kDa) stand-alone MbtH domain was observed, indicating that LtxB is translated as a single polypeptide rather than as separate functional units, at least in *E. coli*.

**Figure 3:**
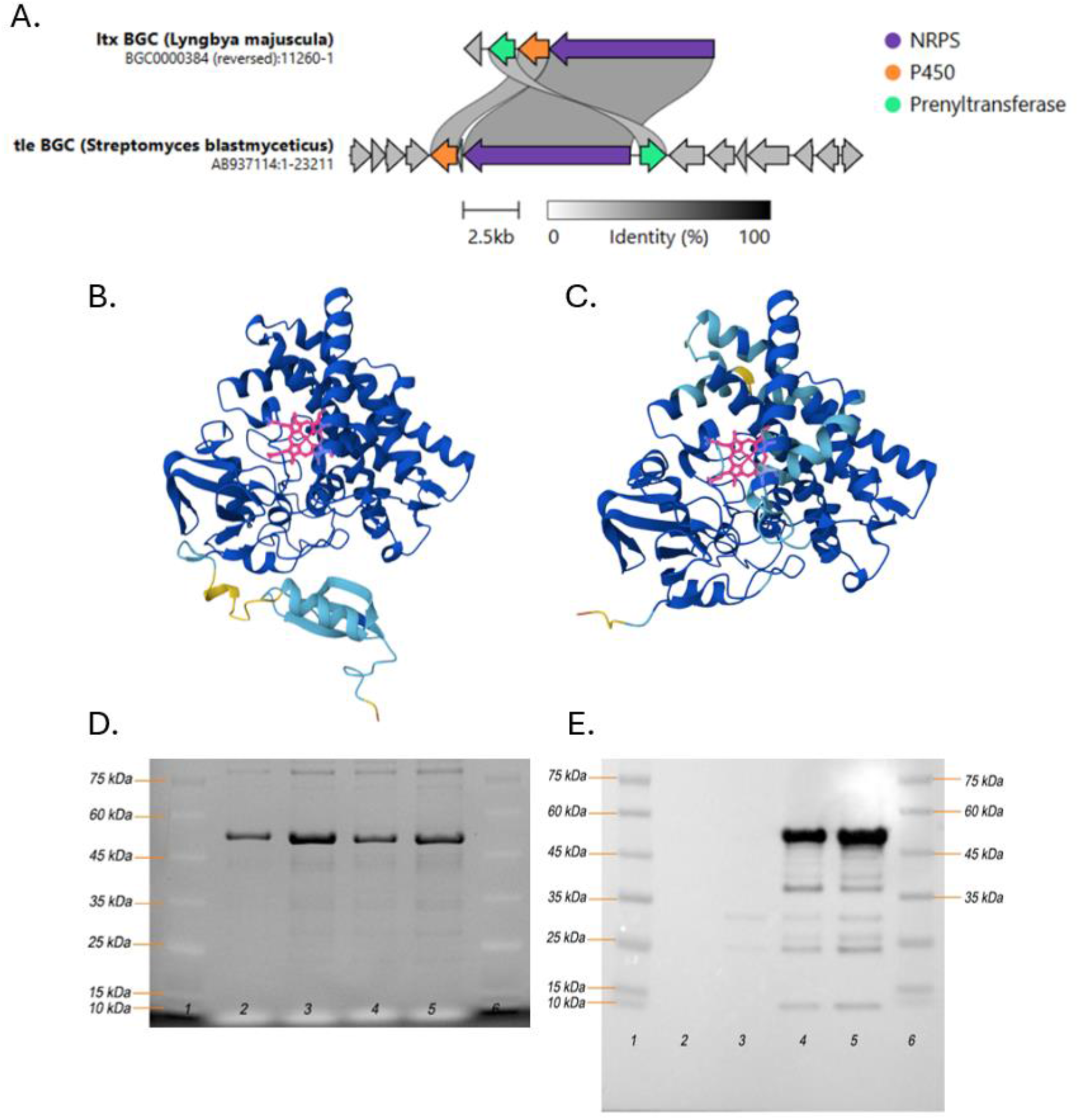
A) Synteny analysis of the *ltx* and *tle* BGCs using clinker; B) AlphaFold model of LtxB with the MbtH domain fusion and the heme cofactor highlighted in pink; C) AlphaFold model of TleB lacking an MbtH domain with heme highlighted in pink; D) SDS-PAGE of recombinant LtxB; E) Western blot of recombinant LtxB using an anti-6xHis tag antibody. In the gel images, lanes 1 and 6 are the protein ladder; 2 and 3 are the first and second eluate of N-terminal 6xHis tagged LtxB; 4 and 5 are first and second eluate of C-terminal 6xHis tagged LtxB.

## Discussion

Heterologous production of cyanobacterial natural products is notoriously challenging in prokaryotic hosts yet the reasons are mostly unknown (Dhakal et al, 2021; Chen et al., 2025). The *ltx* BGC is one of the most well-studied cyanobacterial gene clusters, being heterologously expressed in *E. coli, Anabaena* sp., and *Streptomyces* sp. hosts but has not yet reached industry-relevant yields (Jones et al., 2012; Ongley et al., 2013; Videau et al., 2016). We therefore attempted to heterologously express the *ltx* BGC in *A. oryzae* NSAR1 by cloning individual genes under the control of the strong, inducible *amyB* promoter, essentially over-riding prokaryotic mechanisms for operon expression. Although transcripts could be detected for *ltxA*-*C* and *hetI/sfp*, neither LTXA, ILV, intermediates, analogues, or shunt products were detected. The *ltxA* transcript was truncated, despite the gene being fully integrated into the fungal genome, which appears to be a common phenomenon, as when *ltxA* was heterologously expressed in *Streptomyces coelicolor* A3, a truncated transcript (∼2000 bp) was also reported (Jones et al., 2012). The reasons proposed for truncation include promoter incompatibility, mRNA secondary structure, and / or rare codons, with the latter two proposals also being relevant to our study.

The lack of detectable pathway intermediates in *A. oryzae* may also reflect incompatibility of LtxB with the fungal intracellular environment. LtxB is a bifunctional protein comprising an MbtH-like domain fused to a cytochrome P450 monooxygenase. In contrast to eukaryotic P450 enzymes, which are typically anchored to the endoplasmic reticulum *via* an N-terminal transmembrane helix (Edwards et al., 1991; Lauresen et al., 2021), LtxB lacks such an α-helix. Mislocalization of LtxB in the cytosol may therefore limit access to redox partners required for catalytic activity. In addition, the didomain features of LtxB may present folding challenges in a heterologous host as MbtH-like proteins are often expressed as discrete partners that assist NRPS function (Zwahlen et al., 2019; Madduri et al., 2024). Therefore, fusion to a P450 domain may alter accessibility, folding efficiency, or stability in a eukaryotic system.

Together with the observed truncation of *ltxA* transcripts, these factors highlight multiple barriers to functional reconstitution of cyanobacterial NRPS pathways in fungal hosts. Despite these limitations, successful conversion of ILV to LTXA by codon-optimized LtxC demonstrates that downstream tailoring enzymes can remain functional in *A. oryzae*. This suggests that modular pathway reconstruction offers a viable strategy for engineering cyanobacterial gene clusters in fungal hosts.

## Conclusions

This study demonstrates that full reconstitution of cyanobacterial NRPS pathways in fungal hosts remains challenging due to multiple layers of incompatibility, including transcriptional disruption of large biosynthetic genes and potential mislocalization of associated enzymes. The observed truncation of *ltxA* transcripts highlights transcriptional and splicing barriers as critical limitations when expressing prokaryotic NRPS genes in a eukaryotic system. In addition, the likely incompatibility of LtxB with the fungal intracellular environment further emphasizes the importance of protein localization and domain architecture for pathway functionality.

Importantly, the successful biotransformation of ILV to LTXA by codon-optimized LtxC demonstrates that downstream tailoring enzymes from cyanobacterial pathways can remain functional in *A. oryzae*. These findings support a modular approach to pathway engineering, in which individual enzymes or pathway segments can be selectively reconstructed, and provide opportunities for combinatorial biosynthesis with non-cyanobacterial enzymes. Overall, this work provides practical insight into the limitations and opportunities associated with expressing cyanobacterial biosynthetic pathways in fungal hosts, and highlights key considerations for future efforts to engineer eukaryotic systems for the production and diversification of complex natural products.

## Acknowledgements

This project was supported by UNT Department of Chemistry and BioDiscovery Institute start-up funds (E.J.S), and in part by the W. M. Keck Foundation. We would like to thank Dr. Katherine Williams from UWE Bristol, UK for the pE-YA and pTYGS fungal / yeast / *E. coli* cloning and expression plasmids, as well as the *Aspergillus oryzae* NSAR1 strain. We also want to recognize Jean Christophe Cocuron from the Bioanalytical Facilities (BAF) for acquiring HRMS data. We would like to acknowledge the guidance, protocols, and training received from Emily Herrell. Also, the research technology support facility of Michigan State University and Douglas Whitten for protein sequencing and analysis.

## Author Contributions

SJ constructed plasmids, performed transformations, extractions, feeding studies, LCMS analysis, and protein expression experiments.

TA performed HR-LCMS/MS analysis.

BA and BK performed the timecourse study and prepared organic extracts. CAH and KDC contributed to study design.

ES was responsible for overall study design, supervision of SJ, TA, BA, and BK, funding acquisition, and performed genome mining / bioinformatics analysis, and LCMS analysis.

## Data Availability

