## Supplementary Information for "Heterologous expression of lyngbyatoxin biosynthetic genes in *Aspergillus oryzae* reveals transcriptional barriers but enables LtxC-mediated biotransformation"

### 1. General experimental methods

Analytical grade chemicals and reagents were purchased from Sigma Aldrich and Thermo Fisher Scientific. The solvents used for HPLC were LCMS grade. General molecular biology procedures were performed as standard and molecular biology kits were used according to the manufacturer's protocols. Analytical PCR was performed using OneTaq polymerase and preparative PCR for cloning procedures was performed using Q5 polymerase manufactured by NEB; PCR products were sequenced using the plasmidsaurus sequencing service. Restriction endonucleases were also purchased from NEB. A standard of lyngbyatoxin was purchased from Cayman Chemicals. Synthetic genes (Table S1) and gene fragments (Table S2) were purchased from TWIST Bioscience and resulting plasmids were sequenced using Plasmidsaurus. Oligonucleotides were purchased from IDT (Table S3).

| **Protein Name** | **Amino acid sequence** | **Gene sequence** | **GC %** |
|---|---|---|---|
| LtxA | MIMNQPWSGSRRAIVNSTLPQAFEDRVNRT<br>PNQIAIVFDELHLSYTQLNVKANYLANLLIR<br>SGVEPKTPIAIQIERSIELVVAVLAVLKAGAIY<br>VPLDPRLPLARVHWILDETRAQTLLTISNSK<br>NLDKIKNIQTTIRVDVHLKKEDPDSPAIDTQP<br>DDLAHIMYTSGSTGHPKGVAITHRAILEFVA<br>DRCWKNELQERVLFHSSLGFDISNYELWVP<br>LLRSGQLVMAPSGPLDVSTLKQVIQKQSITS<br>LFLTTSLFNLVTEENPRCLAGVQQVWIGGE<br>QASVAAIQRMWDACPEITVVNGYGPTETTT<br>YVTLYTIESRPQGNRVPIGRPMENTQVYVLD<br>EELKPVPSGTPGEIYLAGTGLARGYFGQPSL<br>TSTRFLANPFGLPGSRMYRTGDLGVWIDSD<br>QLVCLGRSDRQVKIRGIRIEPSEIEAELRNHP<br>EIGEAVVTVREDSPDEKRLIAYLIAKTIESQT<br>QKIQGEQLVAWRQIYDSLYENDTSSPLGSDF<br>SGWHSSYDGCPIPLEQMQEWQEMTLQEIRG<br>LQPRRILEIGVGTGLLLAQLAPDCESYWGT<br>DFSASLIERLRHRVALEPSLAGRVELRSQSA<br>DAIEGLPREHFDTVILNSVIQYFPNPTYLLDV<br>LNKVVDLLVPGGYIFIGDIRNLRLLHCFHTE<br>VQLRRTDLKTLNTLALQAAVEHSVVMEDE<br>LLVDPEFFSALGQRRSDIDYVDVRVKRGWH<br>HNEMSQYRYNVILHKCSTTTEPIPTVEWHPI<br>AWGSKFTTLEQLIDYLKQEIPSHLKLVNVPN<br>CRVTPALKAIQQLNAGYDLDVVQDHLFSGD<br>SAACNPEDFYKIGELLDYQVFVTWSNTDIG<br>HLDILFCPQNSQLNNIRVFRTDSPLHNNQGV<br>YDSLLFYMNSPSVSYHAGTFLESVRTHLSER<br>LSQSMIPTAFVILDTFPLTVNGKVDRAALPV<br>PHIGSTQNGRAPRNAVEERMCGLFAQILGVT<br>QVTIDDNFFDLGGHSLSVMRLLSRIRSCFDV<br>ELAVANVFASPTVAELVKHLTKAEPGRPQLM<br>GMNRPSVVPLSFSQQRLWFIYAFDTSSVIYN<br>VPIAFELAGSINQIAMEMALHDVISRHDSLH<br>TVFQEISGVPRQVLLNPGQFELTITHTSVTEL<br>DQKLLDSSQRPFDLSTEIPIRAELFVLGDDR<br>CVLLILLHHITCDGWSFAPLWRDLSEAYATR<br>SQGLIPDWTPLPVQYADYTLWQRRLLGQDI<br>KPDSIITHQLDYWRKNLAELPELVSLPGDRP<br>RSAVPSFRGALIPFRLSPELHHGAINLAQQQ<br>GTSLFMVLHAVLAVLLTRLGAGTDLPIGTPV<br>ANRTDDALEDLVGFFVNLLVLRTDTSGDPSF | ATGATTATGAATCAACCTTGGAGTGGAAGTAGGCGGGCGATAGTAAATTCGACTTTACCCCAAGCATTTGAG<br>GATCGAGTTAACCGAACTCCAAATCAGATTGCCATCGTTTTTGACGAACTTCACCTGTCCTACACCCAGCTT<br>AATGTTAAAGCTAATTACTTAGCTAATCTCTTAATTCGTTCTGGTGTTGAACCAAAAACTCCTATTGCCATTC<br>AGATAGAGCGTTCGATTGAGTTAGTGGTTGCTGTCCTTGCTGTGCTTAAAGCAGGTGCCATCTATGTTCCTC<br>TAGACCCCCGTCTACCTCTAGCGCGAGTTCATTGGATATTAGATGAAACAAGGGCACAAACATTACTGACTA<br>TAAGTAATTCAAAGAATCTAGACAAGATCAAAAATATTCAAACAACAATTAGAGTAGATGTCCATTTAAAAA<br>AGGAAGATCCAGATAGTCCTGCTATAGATACGCAACCCGATGATCTTGCTCATATTATGTACACCTCAGGCTC<br>GACAGGGCATCCTAAAGGGGTCGCTATTACCCATCGAGCCATTTTAGAATTCGTTGCGGATCGGTGTTGGAA<br>AAATGAACTTCAGGAGCGAGTCCTATTTCACAGTTCTCTAGGCTTTGATATCTCAAATTATGAACTGTGGGT<br>GCCTCTTTTACGATCTGGACAGTTAGTTATGGCCCCTTCTGGACCCTTGGATGTTTCTACTCTTAAACAAGTT<br>ATCCAGAAGCAATCTATTACATCTTTATTCTTAACTACTTCATTATTCAATCTAGTTACTGAGGAAAATCCCAG<br>ATGCCTTGCAGGTGTTCAACAAGTATGGATTGGAGGGGAACAAGCTTCAGTAGCTGCTATTCAGCGGATGT<br>GGGATGCTTGCCCTGAGATTACTGTGGTCAATGGTTATGGTCCTACAGAGACCACCACATATGTCACCCTTT<br>ATACTATTGAAAGTCGCCCTCAGGGCAATCGAGTCCCGATCGGTAGGCCGATGGAAAATACCCAGGTCTAT<br>GTACTTGACGAGGAACTAAAACCCGTTCCTTCCGGTACACCAGGAGAAATCTATCTGGCAGGAACAGGTCT<br>TGCACGAGGCTACTTTGGTCAACCAAGTTTAACATCCACACGGTTCCTTGCTAATCCCTTTGGGCTACCAGG<br>ATCGCGGATGTATCGCACTGGTGATTTAGGAGTGTGGATTGACTCGGATCAGCTTGTCTGTCTTGGTCGCTC<br>TGATCGACAAGTTAAGATCCGTGGCATTCGCATTGAACCTAGCGAAATAGAGGCAGAACTCCGCAATCATC<br>CAGAGATTGGTGAAGCAGTCGTTACAGTTCGCGAGGATTCTCCAGATGAAAAGCGCTTGATTGCCTACCTA<br>ATCGCCAAAACGATTGAATCACAAACGCAGAAAATTCAGGGGGAACAACTGGTAGCTTGGCGTCAAATCT<br>ATGACTCGCTCTACGAAAATGATACTTCTTCTCCTTTGGGATCGGATTTTTCAGGTTGGCACAGTAGCTATGA<br>CGGTTGTCCTATTCCCTTAGAACAGATGCAGGAATGGCAGGAGATGACACTGCAGGAGATTCGTGGGCTTC<br>AGCCTCGGCGAATCTTGGAAATTGGAGTTGGTACTGGTCTGTTGCTAGCTCAATTAGCTCCAGACTGTGAG<br>TCTTATTGGGGAACTGATTTCTCGGCATCTTTAATTGAAAGGTTACGTCATCGGGTTGCCCTTGAGCCATCCC<br>TAGCAGGTCGTGTAGAGCTTCGCTCTCAGAGTGCTGATGCGATCGAAGGGTTACCTCGGGAGCATTTCGAT<br>ACTGTAATTCTCAATTCTGTAATTCAGTATTTTCCTAATCCTACCTATTTGCTTGATGTTCTGAACAAGGTGGT<br>AGATTTGCTTGTTCCAGGTGGTTATATTTTCATTGGTGACATTCGGAACTTGCGTCTTTTGCACTGTTTTCAC<br>ACTGAAGTTCAACTGCGGCGTACTGACCTCAAAACTCTCAACACCCTCGCGTTACAAGCAGCTGTTGAAC<br>ATAGTGTAGTGATGGAAGATGAACTGTTAGTCGATCCTGAGTTTTTTTCAGCTTTAGGTCAAAGACGATCGG<br>ACATTGACTATGTAGATGTGCGAGTTAAAAGGGGTTGGCACCATAATGAAATGTCTCAGTATCGTTACAACG<br>TTATCCTCCACAAATGCTCGACCACAACAGAACCTATACCTACTGTTGAGTGGCATCCGATCGCTTGGGGAT<br>CAAAATTTACAACCCTTGAACAGTTGATTGATTATCTTAAACAGGAGATACCATCCCACCTAAAATTAGTCA<br>ATGTGCCCAACTGTCGCGTTACTCCTGCTCTTAAAGCAATACAACAGTTGAATGCGGGTTACGATCTTGATG<br>TTGTGCAAGATCATCTATTCTCTGGAGACTCTGCTGCTTGCAACCCAGAAGACTTCTATAAGATTGGAGAGT<br>TATTGGATTATCAAGTTTTTGTAACTTGGTCTAATACAGACATAGGCCATCTTGACATCCTGTTTTGTCCCCA<br>GAATAGTCAGTTGAATAACATCAGAGTCTTCAGGACCGATAGTCCTCTTCATAATAATCAGGGGGTATACGA<br>TTCCTTATTGTTTTACATGAACTCTCCCTCTGTCTCATACCATGCAGGAACATTTCTAGAATCAGTGCGGACT<br>CATCTGAGTGAAAGGTTATCTCAGTCGATGATACCCACTGCCTTTGTCATTCTGGATACTTTTCCGTTGACTG<br>TAAATGGCAAAGTCGATCGTGCTGCTTTACCTGTTCCCCATATTGGTTCCACTCAGAATGGACGCGCCCCCA<br>GAAATGCTGTTGAAGAGCGTATGTGTGGCTTGTTCGCTCAAATTCTGGGCGTTACTCAAGTTACAATTGATG<br>ATAATTTTTTTGATCTTGGTGGGCATTCTTTATCTGTGATGCGGTTATTAAGCCGTATCCGCTCATGCTTTGAT<br>GTGGAACTAGCTGTTGCCAATGTTTTTGCTTCACCAACAGTCGCTGAACTGGTCAAACATCTAACCAAAGC<br>AGAACCAGGGCGACCTCAACTCATGGGTATGAACCGCCCCAGTGTTGTACCATTGTCTTTTTCTCAACAAC | 45.4 |

| | | |
|---|---|---|
| | SDLLARIRDLDIAAYAHQDIPFELLVEKLRPV<br>RSITHHPFFEILLALQSAPENQLKLSGVKISH<br>KQVHLGTCRFDLVMNLDEHRHQDGTLLGI<br>DGLVEYRTDLYDRKTIQTFIDRFLRLLTGAIA<br>DPQQRISRLNLLSPEEHHQLLIEWNDTDHSI<br>SPRPFPDLFEAQVAQTPTANALVFGALHLSY<br>QELNARANQLAHQLIYQGIGPGQVIAIDVPR<br>SPEWVIAVLAVLKAGAAYLPLDPSYPASRIIY<br>MLEDVQPCLLITTTNSLISNPKLNIPKLQLDS<br>FPWEASLNVTSEKQDYGLQANIEDNRGTQP<br>LHVSDPAYIIYTSGSTGKPKGVVVTHAGISS<br>MVATQIKYFEVTPESRILQFSSLSFDGVVWE<br>LCSALLTGATLVMAPSERVQPGPELIQLIGD<br>YHVTHAVLPPAVLMVLSPDNIPSLTHLIVSGE<br>AASGELVKRWSVGRCLINGYGPTETTVCAT<br>LSSPLSGNGIPPIGRAVINVQCYVLDDQLQL<br>LPPGAIGELYISGPGLARGYLNQPQLTAERFL<br>ANPFRDIGSRMYRTGDLVRWRNHGELEFVG<br>RADNQVKIRGFRVELGEVETALTNCPPVSD<br>ALAMVREDRPGEKFLVAYVVGQDSMDTDA<br>LRAQLVNDLPPYLIPGAIVTLKKFPLTVNGKI<br>DRKALPVPKFTSIAEGRVPKTPTQIGLCTLFA<br>ELLGVDQVTIDDPFFALGGHSLLATQLVSQI<br>RQRFGISLPVQTVFERQTVADLAIVVDREAP<br>IVSTSIDLAAEVVLNPQIAPYQSRPVELDRN<br>THPASVLLTGATGFLGAYLLYELLKQTDAN<br>VFCLVRSNHSEAAYQRIHSTLKFYQLWSES<br>WRSRIIPVCGDLSQPSLGLSAEEFSKLTELID<br>AIYHNGAQVSAIEPYTYLKPTNVLGTSELLD<br>FAARCRVKPLHFVSTAAVAVSSKGNPDIIYE<br>NFRLGADSVLPSGYVSSKWVAEELVWVASD<br>RGLPVTVHRPGRISGDTTTGIGNTDDTFWQI<br>VRAIVVLGVVPDIVYQDDAGIDLMPVDRVA<br>SAIVHLSRHKQSISKVHHLTCPTIVKLDVVF<br>NELSKLGYQLTTVSYSEWVKQLEQYVDQA<br>PGGHSLASATVLSRTLPKLIELSQICFDQSNT<br>LTGLAEAPFKFPCIDRHLVRGYLTYFINSKFF<br>PQITYTGK* | GTCTGTGGTTTATCTATGCTTTTGATACTTCTAGTGTAATTTATAATGTCCCCATCGCTTTTGAGCTGGCTGGC<br>TCCATCAATCAAATTGCGATGGAGATGGCTTTACATGACGTCATCTCCAGACATGATAGCTTACATACTGTCT<br>TTCAGGAAATTAGCGGTGTACCACGCCAGGTTCTATTGAATCCGGGTCAGTTTGAATTGACTATCACTCACA<br>CTTCTGTAACAGAACTGGATCAGAAACTTCTGGACAGTAGTCAGCGACCTTTTGACCTCAGTACTGAAATT<br>CCAATCCGAGCAGAACTATTTGTCCTGGGAGACGATCGATGTGTCCTCCTCATTCTTCTACACCATATTACCT<br>GCGATGGATGGTCTTTTGCCCCCTTATGGCGCGATCTTTCGGAGGCTTATGCCACCCGTTCCCAGGGTCTCA<br>TCCCTGACTGGACACCGCTACCTGTACAGTATGCAGATTATACCCTATGGCAAAGACGACTCCTTGGCCAAG<br>ACATTAAGCCAGATAGTATTATTACTCATCAACTTGACTATTGGCGCAAAAACCTAGCGGAGCTACCTGAAC<br>TTGTATCCCTTCCTGGAGATCGCCCCCGATCTGCTGTCCCTTCCTTTAGAGGTGCTCTTATTCCCTTTCGGTT<br>GAGCCCTGAACTCCATCATGGGGCGATCAATCTGGCTCAACAACAAGGAACAAGTCTATTCATGGTGTTGC<br>ATGCTGTTCTTGCTGTCCTTTTAACTCGTCTGGGTGCAGGAACCGATCTGCCCATTGGAACCCCTGTGGCTA<br>ATCGCACAGATGATGCACTTGAAGATTTGGTTGGGTTCTTTGTTAACTTGCTCGTTCTACGAACAGATACCT<br>CTGGCGATCCAAGTTTCTCCGATCTCTTAGCCCGAATCAGAGACCTAGATATTGCTGCTTATGCCCATCAAG<br>ATATTCCCTTTGAGTTGTTGGTCGAGAAACTTCGACCCGTGAGATCCATAACTCATCACCCCTTCTTTGAAA<br>TTTTGCTAGCCCTACAGAGTGCACCAGAGAATCAGTTAAAGCTGTCTGGGGTTAAGATAAGTCACAAACAG<br>GTTCATTTGGGAACCTGTCGTTTTGATTTGGTGATGAATCTTGATGAACACCGACATCAGGACGGGACACTC<br>CTGGGAATTGATGGCCTGGTTGAGTATCGTACTGATTTGTACGATCGTAAGACGATTCAGACTTTTATCGATC<br>GCTTCTTGAGACTCTTGACCGGAGCCATTGCTGACCCTCAACAGCGGATTAGTAGACTCAACCTCTTGAGT<br>CCTGAAGAACACCATCAACTCTTGATCGAATGGAATGATACGGACCATAGCATCAGTCCACGACCTTTTCCA<br>GACCTGTTTGAAGCCCAAGTAGCCCAAACTCCTACAGCTAATGCACTTGTATTTGGGGCTTTGCATCTGTCC<br>TATCAAGAACTCAATGCTCGTGCTAACCAACTAGCCCATCAACTGATCTATCAAGGCATAGGTCCAGGCCAA<br>GTCATCGCGATCGATGTTCCACGATCGCCCGAATGGGTCATTGCTGTTTTAGCTGTACTCAAAGCAGGTGCA<br>GCCTACTTACCCCTAGACCCCTCTTACCCAGCCTCTCGCATTATCTATATGCTGGAGGATGTCCAACCCTGCC<br>TGTTAATCACCACAACTAATAGTCTCATTAGTAATCCAAAGCTCAATATTCCAAAACTACAACTCGATAGCTT<br>TCCCTGGGAGGCTTCCCTTAACGTCACCAGTGAAAAACAGGATTATGGATTACAGGCTAATATAGAGGACA<br>ATCGGGGCACTCAGCCACTGCATGTGTCCGATCCAGCCTATATAATTTACACTTCAGGCTCTACAGGAAAAC<br>CCAAAGGAGTTGTGGTAACCCATGCTGGAATTTCGAGTATGGTCGCAACACAAATCAAGTACTTTGAGGTA<br>ACTCCTGAGAGTCGTATCCTCCAGTTCTCTTCTTTAAGTTTTGACGGCGTAGTTTGGGAACTGTGCAGTGCC<br>CTGTTAACTGGTGCCACCTTGGTGATGGCTCCCTCAGAACGGGTGCAACCAGGTCCGGAACTGATCCAATT<br>GATCGGAGACTATCACGTTACCCATGCTGTCTTGCCACCAGCAGTATTAATGGTGCTATCACCAGATAATATA<br>CCCTCCTTGACCCATCTCATTGTCTCAGGCGAGGCTGCCTCAGGTGAATTAGTAAAACGGTGGTCTGTTGG<br>GCGCTGCTTGATTAATGGTTATGGTCCCACTGAAACTACCGTCTGTGCTACCTTGAGTTCTCCCCTATCTGGT<br>AATGGGATACCCCCCATTGGCAGAGCGGTTATCAACGTTCAATGTTATGTCCTTGATGATCAATTGCAACTTC<br>TTCCCCCTGGAGCGATCGGTGAACTTTACATATCTGGCCCTGGCCTAGCCCGTGGATATCTGAACCAGCCTC<br>AGCTTACGGCAGAGCGATTCTTGGCAAATCCCTTTAGGGACATCGGAAGCCGTATGTATCGGACTGGAGAT<br>TTAGTTCGTTGGCGAAACCACGGTGAGCTTGAGTTTGTAGGCAGAGCTGATAATCAGGTCAAGATTCGGGG<br>TTTTCGTGTTGAATTGGGTGAAGTAGAAACTGCCCTTACCAATTGTCCCCCAGTGTCAGATGCCCTCGCTAT<br>GGTTCGTGAGGATCGACCGGGCGAAAAATTCCTAGTGGCCTATGTGGTTGGTCAGGATTCTATGGATACTGA<br>TGCTCTGCGTGCCCAGCTCGTTAATGATTTACCTCCATACTTGATCCCTGGCGCTATTGTCACTTTGAAGAAA<br>TTTCCCCTGACAGTGAACGGTAAAATAGATCGTAAGGCTCTCCCAGTTCCAAAGTTCACTTCTATTGCTGAA<br>GGGCGTGTTCCCAAAACTCCTACCCAGATTGGTTTATGTACTCTGTTTGCAGAACTGTTGGGTGTTGATCAA<br>GTCACGATCGACGATCCGTTTTTTGCCCTTGGAGGTCACTCGTTGCTAGCCACACAATTAGTGAGTCAGATT<br>CGGCAGCGTTTTGGGATAAGTTTGCCAGTGCAAACTGTATTCGAGCGACAGACGGTGGCTGACCTGGCAAT<br>AGTAGTAGATCGGGAAGCACCTATAGTTTCTACTTCCATTGACCTTGCTGCTGAAGTTGTTCTTAATCCGCA<br>GATTGCTCCCTACCAGAGTCGTCCAGTTGAACTCGATAGGAATACTCACCCAGCCTCGGTGTTATTAACCGG<br>AGCAACTGGCTTTTTGGGGGCATATTTGCTTTATGAACTTCTGAAGCAGACTGATGCTAATGTATTTTGCTTG<br>GTGCGCTCCAACCACTCTGAAGCTGCCTATCAACGCATCCATTCGACTCTCAAATTTTATCAATTATGGTCTG<br>AATCTTGGAGATCGCGAATCATTCCAGTGTGTGGCGATCTCTCCCAGCCAAGTTTAGGCTTATCCGCTGAGG<br>AATTCAGCAAGCTCACAGAATTAATTGATGCTATCTATCACAATGGAGCCCAGGTCAGTGCTATTGAACCCT<br>ATACATACCTTAAACCAACAAATGTATTGGGGACATCAGAGTTACTGGACTTTGCTGCGCGATGCCGCGTGA |

| | | | |
|---|---|---|---|
| | | AGCCATTGCACTTTGTTTCAACAGCTGCAGTGGCTGTAAGTTCTAAAGGTAATCCTGATATAATTTATGAGA<br>ATTTTAGACTTGGAGCAGATTCTGTATTACCCAGTGGATATGTCTCTAGTAAATGGGTCGCTGAAGAGTTAG<br>TGTGGGTGGCATCAGACCGTGGCCTCCCCGTTACTGTCCACCGCCCAGGCCGAATTAGTGGTGATACCACT<br>ACTGGTATTGGGAATACTGATGATACATTCTGGCAAATAGTCAGGGCAATAGTTGTTCTAGGGGTTGTCCCA<br>GATATTGTTTATCAAGATGATGCCGGGATCGATTTAATGCCAGTCGATCGCGTTGCAAGTGCAATTGTCCATC<br>TATCCCGTCATAAACAATCCATAAGCAAGGTGCATCACCTCACCTGTCCTACAATAGTCAAGCTAGATGTAG<br>TGTTTAATGAACTCTCTAAATTGGGCTACCAGTTAACCACGGTATCCTACTCTGAATGGGTTAAGCAATTAG<br>AACAGTACGTCGATCAAGCCCCAGGGGGGCACTCCCTAGCATCAGCTACCGTCCTTAGTCGTACATTACCC<br>AAGCTCATAGAACTCAGTCAAATCTGCTTCGACCAAAGCAACACCTTAACGGGTTTAGCTGAAGCACCATT<br>TAAGTTTCCTTGTATTGATCGTCATCTTGTCCGTGGATACCTCACTTATTTCATCAACTCTAAATTCTTCCCTC<br>AAATAACTTATACAGGAAAGTAA | |
| LtxB | MTNPFADPSRDYWVLCNAEGQYSLWPTSL<br>EIPEGWQTAFGPESWQNCLDNVEKNWTDM<br>RPLSLLGQRKCPVPYPFREYVALNMDPIYAS<br>LQQRDEMLRISVPYGDDAWLASSYKHVKC<br>VIQDTRFSRETTKYDEARLTPIPIRTSVLGMD<br>PPDHTRLREALAAALTPSHVEQMKPWITSM<br>TEQLIDDFVAQGPPADLVEQFALPFTGHITCE<br>LMGIPLEDRPQFKAWCDGFSSTSSLTKEEVE<br>VRMQAMYTYITELVGRRRESPSDDIISKLLQ<br>PEDKRLELSEIELIDLATILLLAGYDSTAMEL<br>ANAIFVLLTHPEQTQLIREQPKLMPQAVQEL<br>LRYIPIDAHVTFARYATEDVQVGNTLVRTGD<br>AILASFPSANHDPAIFEDPHTFNIMRPRKPNF<br>GLGTGIHSCAGKLLAVVELEIALSVLIRRLPT<br>LRLAIPAEEVPWQPGSLLRSTSKLPAEW | ATGACAAATCCTTTTGCAGACCCTTCAAGGGACTACTGGGTACTATGCAATGCCGAAGGTCAATATTCCCTA<br>TGGCCCACTTCCCTTGAAATTCCTGAAGGCTGGCAAACCGCTTTCGGCCCCGAGAGTTGGCAAAATTGCCT<br>TGACAACGTTGAGAAAAACTGGACTGACATGCGCCCCCTCAGTCTGTTAGGTCAACGGAAGTGTCCTGTG<br>CCTTACCCCTTTCGAGAATATGTTGCCCTTAACATGGATCCAATCTACGCCAGTCTGCAACAACGGGATGAA<br>ATGCTACGAATTAGTGTTCCCTATGGGGATGATGCCTGGTTAGCCTCCAGTTACAAACATGTCAAGTGTGTT<br>ATACAAGATACACGTTTTAGCCGAGAAACAACTAAGTATGATGAGGCTCGCTTAACACCAATACCGATCCGC<br>ACGAGTGTATTGGGGATGGATCCTCCAGACCATACACGATTACGTGAGGCTTTAGCCGCAGCTCTCACTCCT<br>AGTCATGTTGAGCAAATGAAGCCTTGGATAACCAGCATGACAGAACAACTGATCGATGACTTTGTGGCCCA<br>AGGTCCCCCTGCCGATCTAGTCGAGCAATTTGCTTTGCCATTTACTGGACACATAACTTGCGAACTCATGGG<br>AATTCCGCTAGAAGATCGTCCCCAATTCAAAGCATGGTGTGATGGATTTTCCTCCACTTCGTCTCTCACCAA<br>AGAGGAAGTAGAAGTTCGTATGCAGGCAATGTATACCTATATTACTGAGCTTGTGGGGCGACGACGGGAAT<br>CTCCAAGTGATGACATTATTAGTAAATTACTGCAACCTGAGGATAAGCGCCTGGAGCTATCAGAAATTGAAC<br>TGATTGACCTAGCCACGATCTTATTACTTGCAGGATATGACTCAACAGCTATGGAACTTGCAAATGCGATTTT<br>TGTACTTTTAACCCACCCAGAACAGACCCAACTCATCCGAGAACAACCCAAGCTGATGCCCCAAGCTGTGC<br>AAGAATTGCTCCGCTATATTCCCATTGACGCCCATGTTACTTTTGCTCGCTATGCCACTGAGGACGTTCAGGT<br>TGGTAATACGCTTGTTCGCACTGGTGATGCAATTTTGGCGTCTTTCCCCTCTGCCAATCACGATCCTGCAATA<br>TTTGAAGATCCTCACACATTCAATATCATGCGACCTCGCAAACCAAACTTTGGCTTAGGAACTGGTATACAC<br>AGTTGTGCTGGTAAGCTCCTGGCTGTTGTGGAGTTGGAGATTGCACTATCTGTCTTGATTAGACGTCTGCCT<br>ACTTTACGTTTAGCTATTCCAGCTGAAGAAGTGCCATGGCAACCAGGCTCTCTGTTGCGCAGTACTTCAAA<br>GTTACCTGCTGAGTGGTAA | 46.9 |
| LtxC | MNSKIAVSEYASDSFGKADTRAYDDLPIFAS<br>NYYSREASYIDIACDKLHRACDGLAFPAER<br>RDTIVAFLRKLCTPWGNALASAPQHFSNVS<br>VEGMPFELSLAWSNGRGHELRMSFEVLHSP<br>HSLSNNLEAARRFVRELPSIPELAPDISIENFL<br>LIEDLVTASEPTDFITPMGQGIAWLSDRPNPV<br>MKTYLNLNIAGFESGPERTKEAMTRLGAGS<br>AWQSLCSYLTSLGVSVVPLFLALDLAHSDH<br>PRLKIYLPHTGVDAGMIDKQAEIAQTHVNG<br>KFKEALLEITGHTTPDWRKVPVTCYTLVPG<br>QDRPVAATLYVPLNPNIENDAVAQERVCAFS<br>QSQGVDPSPYESLLKAISDKPLSQLSTHNFV<br>AYRPGKDPRFSVYLAPGVYKRS | ATGAATTCAAAGATCGCTGTTTCTGAATACGCCAGTGATAGTTTTGGTAAGGCTGATACTCGGGCCTATGAT<br>GACCTGCCAATTTTCGCTTCAAATTATTACTCACGAGAAGCGAGTTATATTGATATTGCCTGTGACAAACTAC<br>ATCGAGCGTGTGATGGATTGGCCTTCCCAGCAGAGCGTAGGGATACCATAGTTGCCTTCCTTCGTAAACTCT<br>GTACCCCATGGGGTAACGCTTTAGCTAGTGCTCCGCAGCATTTTTCTAATGTATCTGTTGAGGGAATGCCCTT<br>TGAACTATCCCTGGCCTGGAGCAATGGTCGAGGTCATGAATTACGGATGTCTTTTGAAGTCCTCCATAGCCC<br>CCATAGCTTGAGTAATAATCTTGAAGCTGCCCGTCGATTTGTGCGAGAGCTGCCCAGTATTCCTGAATTGGC<br>TCCGGATATCTCCATTGAGAACTTTCTTCTAATTGAAGATTTAGTGACTGCTTCCGAGCCAACAGATTTTATT<br>ACTCCCATGGGTCAGGGTATTGCTTGGTTATCTGACAGACCAAATCCTGTGATGAAAACCTACCTTAATCTC<br>AATATTGCTGGTTTTGAATCTGGGCCTGAGCGAACTAAGGAAGCCATGACTAGGTTAGGTGCAGGTTCTGC<br>CTGGCAATCACTTTGTTCCTATCTCACTAGCCTGGGAGTGTCGGTTGTTCCCCTCTTCCTAGCTCTCGATCTG<br>GCTCATTCAGACCATCCCAGACTCAAGATTTATTTACCCCATACAGGGGTTGATGCTGGGATGATTGATAAA<br>CAGGCAGAAATTGCTCAAACCCATGTTAACGGGAAATTTAAAGAGGCATTACTAGAGATTACTGGGCATAC<br>AACACCAGACTGGCGAAAAGTCCCTGTTACTTGCTATACCCTCGTTCCAGGGCAAGATCGCCCCGTGGCAG<br>CAACCCTTTATGTTCCGCTCAATCCCAATATTGAAAATGATGCAGTCGCCCAAGAACGGGTCTGTGCCTTCT<br>CCCAATCTCAAGGAGTTGACCCCTCACCCTACGAGTCATTACTCAAGGCAATCTCCGATAAGCCACTATCTC<br>AATTATCAACCCACAATTTCGTAGCCTATCGACCAGGCAAAGATCCGCGCTTTTCAGTTTATCTTGCACCAG<br>GAGTCTATAAACGGTCATAA | 46.1 |
| LtxC-co | MNSKIAVSEYASDSFGKADTRAYDDLPIFAS<br>NYYSREASYIDIACDKLHRACDGLAFPAER | ATGAACAGCAAAATTGCTGTATCCGAGTATGCCTCGGATAGTTTCGGCAAAGCTGATACACGCGCCTATGAC<br>GACCTTCCAATCTTCGCCAGCAATTATTACTCCCGCGAAGCTTCCTATATCGATATTGCCTGCGACAAACTCC | 48.1 |

| | | | |
|---|---|---|---|
| | RDTIVAFLRKLCTPWGNALASAPQHFSNVS<br>VEGMPFELSLAWSNGRGHELRMSFEVLHSP<br>HSLSNNLEAARRFVRELPSIPELAPDISIENFL<br>LIEDLVTASEPTDFITPMGQGIAWLSDRPNPV<br>MKTYLNLNIAGFESGPERTKEAMTRLGAGS<br>AWQSLCSYLTSLGVSVVPLFLALDLAHSDH<br>PRLKIYLPHTGVDAGMIDKQAEIAQTHVNG<br>KFKEALLEITGHTTPDWRKVPVTCYTLVPG<br>QDRPVAATLYVPLNPNIENDAVAQERVCAFS<br>QSQGVDPSPYESLLKAISDKPLSQLSTHNFV<br>AYRPGKDPRFSVYLAPGVYKRS | ATCGCGCCTGCGATGGCTTGGCTTTCCCAGCGGAGAGACGTGATACTATCGTGGCGTTTCTGCGTAAGCTGT<br>GCACTCCCTGGGGAAATGCCCTTGCAAGCGCCCCGCAACACTTTAGCAATGTTTCCGTGGAGGGCATGCCT<br>TTCGAATTGTCCCTCGCTTGGTCGAATGGCAGAGGACATGAGCTTCGTATGTCCTTTGAGGTCTTGCACTCC<br>CCGCATTCCCTTAGCAATAACCTGGAGGCAGCGCGCCGGTTTGTGCGCGAATTGCCATCAATTCCTGAGTTG<br>GCGCCGGACATCTCTATAGAGAATTTCTTGCTGATTGAGGACCTGGTTACGGCTTCTGAGCCCACAGATTTC<br>ATCACTCCGATGGGACAAGGTATCGCGTGGCTTTCTGATCGACCCAACCCCGTAATGAAGACCTATCTGAA<br>CCTTAACATTGCTGGTTTCGAGAGCGGTCCCGAGCGGACCAAGGAGGCCATGACGCGATTGGGTGCGGGA<br>AGCGCTTGGCAATCTCTTTGCTCATACTTGACATCACTGGGAGTGTCCGTTGTGCCCTTGTTTCTCGCACTG<br>GACCTGGCTCATTCAGACCATCCACGTTTGAAGATCTACCTTCCGCACACCGGCGTTGACGCAGGAATGAT<br>CGACAAACAGGCCGAGATAGCTCAGACTCACGTTAACGGCAAATTCAAAGAGGCGCTCCTGGAAATCACA<br>GGTCACACTACACCTGATTGGAGGAAAGTCCCCGTTACATGCTACACTCTCGTTCCCGGCCAAGATAGGCC<br>CGTTGCAGCAACACTCTATGTCCCGCTCAATCCAAACATTGAAAACGATGCGGTGGCGCAAGAGCGGGTTT<br>GCGCTTTCTCGCAATCCCAAGGTGTGGATCCATCGCCTTACGAGTCACTGCTCAAGGCAATTTCCGATAAAC<br>CGTTGAGTCAACTCTCCACCCACAATTTCGTGGCCTATCGGCCTGGTAAGGATCCCAGATTCTCAGTGTATC<br>TTGCTCCCGGTGTATACAAACGGTCTTAA | |
| HetI | MLQHTWLPKPPNLTLLSDEVHLWRIPLDQP<br>ESQLQDLAATLSSDELARANRFYFPEHRRRF<br>TAGRGILRSILGGYLGVEPGQVKFDYESRG<br>KPILGDRFAESGLLFNLSHSQNLALCAVNYT<br>RQIGIDLEYLRPTSDLESLAKRFFLPREYELL<br>RSLPDEQKQKIFFRYWTCKEAYLKATGDGI<br>AKLEEIEIALTPTEPAKLQTAPAWSLLELVPD<br>DNCVAAVAVAGFGWQPKFWHY* | ATGCTGCAACACACCTGGCTCCCGAAACCTCCGAATCTCACCCTGCTTAGCGATGAAGTGCATCTCTGGAG<br>GATCCCCCTAGATCAACCTGAATCTCAACTTCAGGATCTGGCCGCTACCCTAAGCTCGGATGAACTCGCCCG<br>CGCAAACAGGTTCTATTTCCCAGAGCATCGAAGGCGATTCACTGCCGGACGAGGCATTCTCAGGTCCATCC<br>TAGGTGGCTACCTCGGAGTCGAGCCAGGACAGGTCAAGTTCGATTACGAGAGTCGGGGTAAACCAATCCT<br>GGGTGATAGGTTCGCTGAATCCGGACTTCTCTTCAACCTTAGCCACAGCCAGAACCTCGCCTTGTGCGCTG<br>TTAATTACACCCGGCAGATAGGAATCGATCTCGAATATCTGCGCCCGACTAGCGATTTGGAATCCCTGGCTA<br>AGCGATTCTTCCTACCTCGGGAGTACGAGCTGTTGAGGAGCCTGCCCGATGAACAGAAACAAAAGATTTTC<br>TTCCGGTACTGGACGTGCAAGGAGGCTTACCTTAAGGCAACTGGCGATGGCATCGCCAAGCTCGAAGAGA<br>TCGAGATCGCACTGACCCCGACCGAGCCAGCGAAGCTCCAGACAGCCCCAGCCTGGTCCCTGCTGGAGTT<br>GGTCCCCGATGACAACTGTGTAGCTGCTGTTGCTGTGGCCGGTTTTGGCTGGCAGCCCAAATTCTGGCACT<br>ATTAA | 54.9 |
| Sfp | MKIYGIYMDRPLSQEENERFMSFISPEKREK<br>CRRFYHKEDAHRTLLGDVLVRSVISRQYQL<br>DKSDIRFSTQEYGKPCIPDLPDAHFNISHSGR<br>WVICAFDSQPIGIDIEKTKPISLEIAKRFFSKT<br>EYSDLLAKDKDEQTDYFYHLWSMKESFIKQ<br>EGKGLSLPLDSFSVRLHQDGQVSIELPDSHS<br>PCYIKTYEVDPGYKMAVCAAHPDFPEDITM<br>VSYEELL* | ATGAAGATCTACGGAATATACATGGATCGTCCGCTCTCGCAAGAGGAAAACGAGCGCTTCATGTCCTTTATC<br>AGTCCAGAGAAACGTGAAAAATGCAGACGGTTCTACCACAAGGAGGACGCACACCGAACCCTGCTGGGA<br>GATGTCTTGGTTCGTTCCGTCATCTCTAGGCAATATCAACTAGACAAGTCAGATATCCGTTTTTCCACTCAAG<br>AGTACGGCAAACCGTGCATTCCCGACTTGCCCGACGCCCACTTCAACATCAGCCACAGTGGCCGATGGGTT<br>ATATGCGCTTTTGACTCTCAACCAATTGGAATTGATATTGAAAAGACGAAACCGATATCGCTCGAAATTGCC<br>AAACGCTTTTTCAGTAAGACGGAGTATTCTGACTTGCTTGCAAAGGATAAGGACGAGCAGACGGATTATTT<br>TTACCATCTGTGGAGTATGAAGGAATCCTTTATAAAACAGGAGGGAAAAGGTCTTTCCCTTCCTTTGGACTC<br>ATTTTCCGTGAGGCTTCATCAAGATGGACAGGTATCAATAGAGCTCCCCGACTCACATTCACCCTGCTACAT<br>CAAGACTTACGAGGTTGATCCTGGTTACAAAATGGCAGTCTGTGCGGCTCATCCAGACTTTCCTGAAGATAT<br>CACAATGGTTTCTTACGAGGAGTTGCTGTAG | 46.0 |

**Table S1**: Synthetic genes used in this study with start and stop codons underlined.

| Gene fragment name | Sequence |
|---|---|
| LtxA-F1 | AATGCCAACTTTGTACAAAAAAGCAGGCTCATGATTATGAATCAACCTTGGAGTGGAAGTAGGCGGGCGATAGTAAATTCGACTTTACCCCAAGCATTTGAGGATCGA<br>GTTAACCGAACTCCAAATCAGATTGCCATCGTTTTTGACGAACTTCACCTGTCCTACACCCAGCTTAATGTTAAAGCTAATTACTTAGCTAATCTCTTAATTCGTTCTGGT<br>GTTGAACCAAAAACTCCTATTGCCATTCAGATAGAGCGTTCGATTGAGTTAGTGGTTGCTGTCCTTGCTGTGCTTAAAGCAGGTGCCATCTATGTTCCTCTAGACCCCC<br>GTCTACCTCTAGCGCGAGTTCATTGGATATTAGATGAAACAAGGGCACAAACATTACTGACTATAAGTAATTCAAAGAATCTAGACAAGATCAAAAATATTCAAACAAC |

| | |
|---|---|
| | AATTAGAGTAGATGTCCATTTAAAAAAGGAAGATCCAGATAGTCCTGCTATAGATACGCAACCCGATGATCTTGCTCATATTATGTACACCTCAGGCTCGACAGGGCATC<br>CTAAAGGGGTCGCTATTACCCATCGAGCCATTTTAGAATTCGTTGCGGATCGGTGTTGGAAAAATGAACTTCAGGAGCGAGTCCTATTTCACAGTTCTCTAGGCTTTGA<br>TATCTCAAATTATGAACTGTGGGTGCCTCTTTTACGATCTGGACAGTTAGTTATGGCCCCTTCTGGACCCTTGGATGTTTCTACTCTTAAACAAGTTATCCAGAAGCAAT<br>CTATTACATCTTTATTCTTAACTACTTCATTATTCAATCTAGTTACTGAGGAAAATCCCAGATGCCTTGCAGGTGTTCAACAAGTATGGATTGGAGGGGAACAAGCTTCA<br>GTAGCTGCTATTCAGCGGATGTGGGATGCTTGCCCTGAGATTACTGTGGTCAATGGTTATGGTCCTACAGAGACCACCACATATGTCACCCTTTATACTATTGAAAGTCG<br>CCCTCAGGGCAATCGAGTCCCGATCGGTAGGCCGATGGAAAATACCCAGGTCTATGTACTTGACGAGGAACTAAAACCCGTTCCTTCCGGTACACCAGGAGAAATCTA<br>TCTGGCAGGAACAGGTCTTGCACGAGGCTACTTTGGTCAACCAAGTTTAACATCCACACGGTTCCTTGCTAATCCCTTTGGGCTACCAGGATCGCGGATGTATCGCACT<br>GGTGATTTAGGAGTGTGGATTGACTCGGATCAGCTTGTCTGTCTTGGTCGCTCTGATCGACAAGTTAAGATCCGTGGCATTCGCATTGAACCTAGCGAAATAGAGGCA<br>GAACTCCGCAATCATCCAGAGATTGGTGAAGCAGTCGTTACAGTTCGCGAGGATTCTCCAGATGAAAAGCGCTTGATTGCCTACCTAATCGCCAAAACGATTGAATCA<br>CAAACGCAGAAAATTCAGGGGGAACAACTGGTAGCTTGGCGTCAAATCTATGACTCGCTCTACGAAAATGATACTTCTTCT |
| LtxA-F2 | GACTCGCTCTACGAAAATGATACTTCTTCTCCTTTGGGATCGGATTTTTCAGGTTGGCACAGTAGCTATGACGGTTGTCCTATTCCCTTAGAACAGATGCAGGAATGGC<br>AGGAGATGACACTGCAGGAGATTCGTGGGCTTCAGCCTCGGCGAATCTTGGAAATTGGAGTTGGTACTGGTCTGTTGCTAGCTCAATTAGCTCCAGACTGTGAGTCTT<br>ATTGGGGAACTGATTTCTCGGCATCTTTAATTGAAAGGTTACGTCATCGGGTTGCCCTTGAGCCATCCCTAGCAGGTCGTGTAGAGCTTCGCTCTCAGAGTGCTGATGC<br>GATCGAAGGGTTACCTCGGGAGCATTTCGATACTGTAATTCTCAATTCTGTAATTCAGTATTTCCTAATCCTACCTATTTGCTTGATGTTCTGAACAAGGTGGTAGATTT<br>GCTTGTTCCAGGTGGTTATATTTTCATTGGTGACATTCGGAACTTGCGTCTTTTGCACTGTTTTCACACTGAAGTTCAACTGCGGCGTACTGACCTCAAAACTCTCAAC<br>ACCCTCGCGTTACAAGCAGCTGTTGAACATAGTGTAGTGATGGAAGATGAACTGTTAGTCGATCCTGAGTTTTTTTCAGCTTTAGGTCAAAGACGATCGGACATTGACT<br>ATGTAGATGTGCGAGTTAAAAGGGGTTGGCACCATAATGAAATGTCTCAGTATCGTTACAACGTTATCCTCCACAAATGCTCGACCACAACAGAACCTATACCTACTGT<br>TGAGTGGCATCCGATCGCTTGGGGATCAAAATTTACAACCCTTGAACAGTTGATTGATTATCTTAAACAGGAGATACCATCCCACCTAAAATTAGTCAATGTGCCCAAC<br>TGTCGCGTTACTCCTGCTCTTAAAGCAATACAACAGTTGAATGCGGGTTACGATCTTGATGTTGTGCAAGATCATCTATTCTCTGGAGACTCTGCTGCTTGCAACCCAG<br>AAGACTTCTATAAGATTGGAGAGTTATTGGATTATCAAGTTTTTGTAACTTGGTCTAATACAGACATAGGCCATCTTGACATCCTGTTTTGTCCCCAGAATAGTCAGTTG<br>AATAACATCAGAGTCTTCAGGACCGATAGTCCTCTTCATAATAATCAGGGGGTATACGATTCCTTATTGTTTTACATGAACTCTCCCTCTGTCTCATACCATGCAGGAACA<br>TTTCTAGAATCAGTGCGGACTCATCTGAGTGAAAGGTTATCTCAGTCGATGATACCCACTGCCTTTGTCATTCTGGATACTTTTCCGTTGACTGTAAATGGCAAAGTCGA<br>TCGTGCTGCTTTACCTGTTCCCCATATTGGTTCCACTCAGAATGGACGCGCCCCCAGAAATGCTGTTGAAGAGCGTATGTGTGGCTTGTTCGCTCAAATTCTGGGCGTT<br>ACTCAAGTTACAATTGATGATAATTTTTTTGATCTTGGTGGGCATTCTTTATCTGTGATGCGGTTATTAAGCCGTATC |
| LtxA-F3 | TTATCTGTGATGCGGTTATTAAGCCGTATCCGCTCATGCTTTGATGTGGAACTAGCTGTTGCCAATGTTTTTGCTTCACCAACAGTCGCTGAACTGGTCAAACATCTAAC<br>CAAAGCAGAACCAGGGCGACCTCAACTCATGGGTATGAACCGCCCCAGTGTTGTACCATTGTCTTTTTCTCAACAACGTCTGTGGTTTATCTATGCTTTTGATACTTCTA<br>GTGTAATTTATAATGTCCCCATCGCTTTTGAGCTGGCTGGCTCCATCAATCAAATTGCGATGGAGATGGCTTTACATGACGTCATCTCCAGACATGATAGCTTACATACTG<br>TCTTTCAGGAAATTAGCGGTGTACCACGCCAGGTTCTATTGAATCCGGGTCAGTTTGAATTGACTATCACTCACACTTCTGTAACAGAACTGGATCAGAAACTTCTGGA<br>CAGTAGTCAGCGACCTTTTGACCTCAGTACTGAAATTCCAATCCGAGCAGAACTATTTGTCCTGGGAGACGATCGATGTGTCCTCCTCATTCTTCTACACCATATTACCT<br>GCGATGGATGGTCTTTTGCCCCCTTATGGCGCGATCTTTCGGAGGCTTATGCCACCCGTTCCCAGGGTCTCATCCCTGACTGGACACCGCTACCTGTACAGTATGCAGA<br>TTATACCCTATGGCAAAGACGACTCCTTGGCCAAGACATTAAGCCAGATAGTATTATTACTCATCAACTTGACTATTGGCGCAAAAACCTAGCGGAGCTACCTGAACTT<br>GTATCCCTTCCTGGAGATCGCCCCCGATCTGCTGTCCCTTCCTTTAGAGGTGCTCTTATTCCCTTTCGGTTGAGCCCTGAACTCCATCATGGGGCGATCAATCTGGCTCA<br>ACAACAAGGAACAAGTCTATTCATGGTGTTGCATGCTGTTCTTGCTGTCCTTTTAACTCGTCTGGGTGCAGGAACCGATCTGCCCATTGGAACCCCTGTGGCTAATCGC<br>ACAGATGATGCACTTGAAGATTTGGTTGGGTTCTTTGTTAACTTGCTCGTTCTACGAACAGATACCTCTGGCGATCCAAGTTTCTCCGATCTCTTAGCCCGAATCAGAG<br>ACCTAGATATTGCTGCTTATGCCCATCAAGATATTCCCTTTGAGTTGTTGGTCGAGAAACTTCGACCCGTGAGATCCATAACTCATCACCCCTTCTTTGAAATTTTGCTA<br>GCCCTACAGAGTGCACCAGAGAATCAGTTAAAGCTGTCTGGGGTTAAGATAAGTCACAAACAGGTTCATTTGGGAACCTGTCGTTTTGATTTGGTGATGAATCTTGAT<br>GAACACCGACATCAGGACGGGACACTCCTGGGAATTGATGGCCTGGTTGAGTATCGTACTGATTTGTACGATCGTAAGACGATTCAGACTTTTATCGATCGCTTCTTGA<br>GACTCTTGACCGGAGCCATTGCTGACCCTCAACAGCGGATTAGTAGACTCAACCTCTTGAGTCCTGAAGAACACCAT |
| LtxA-F4 | CTCAACCTCTTGAGTCCTGAAGAACACCATCAACTCTTGATCGAATGGAATGATACGGACCATAGCATCAGTCCACGACCTTTTCCAGACCTGTTTGAAGCCCAAGTA<br>GCCCAAACTCCTACAGCTAATGCACTTGTATTTGGGGCTTTGCATCTGTCCTATCAAGAACTCAATGCTCGTGCTAACCAACTAGCCCATCAACTGATCTATCAAGGCAT<br>AGGTCCAGGCCAAGTCATCGCGATCGATGTTCCACGATCGCCCGAATGGGTCATTGCTGTTTTAGCTGTACTCAAAGCAGGTGCAGCCTACTTACCCCTAGACCCCTCT<br>TACCCAGCCTCTCGCATTATCTATATGCTGGAGGATGTCCAACCCTGCCTGTTAATCACCACAACTAATAGTCTCATTAGTAATCCAAAGCTCAATATTCCAAAACTACA<br>ACTCGATAGCTTTCCCTGGGAGGCTTCCCTTAACGTCACCAGTGAAAAACAGGATTATGGATTACAGGCTAATATAGAGGACAATCGGGGCACTCAGCCACTGCATGT<br>GTCCGATCCAGCCTATATAATTTACACTTCAGGCTCTACAGGAAAACCCAAAGGAGTTGTGGTAACCCATGCTGGAATTTCGAGTATGGTCGCAACACAAATCAAGTAC<br>TTTGAGGTAACTCCTGAGAGTCGTATCCTCCAGTTCTCTTCTTTAAGTTTTGACGGCGTAGTTTGGGAACTGTGCAGTGCCCTGTTAACTGGTGCCACCTTGGTGATGG<br>CTCCCTCAGAACGGGTGCAACCAGGTCCGGAACTGATCCAATTGATCGGAGACTATCACGTTACCCATGCTGTCTTGCCACCAGCAGTATTAATGGTGCTATCACCAGA<br>TAATATACCCTCCTTGACCCATCTCATTGTCTCAGGCGAGGCTGCCTCAGGTGAATTAGTAAAACGGTGGTCTGTTGGGCGCTGCTTGATTAATGGTTATGGTCCCACTG<br>AAACTACCGTCTGTGCTACCTTGAGTTCTCCCCTATCTGGTAATGGGATACCCCCCATTGGCAGAGCGGTTATCAACGTTCAATGTTATGTCCTTGATGATCAATTGCAA<br>CTTCTTCCCCCTGGAGCGATCGGTGAACTTTACATATCTGGCCCTGGCCTAGCCCGTGGATATCTGAACCAGCCTCAGCTTACGGCAGAGCGATTCTTGGCAAATCCCT<br>TTAGGGACATCGGAAGCCGTATGTATCGGACTGGAGATTTAGTTCGTTGGCGAAACCACGGTGAGCTTGAGTTTGTAGGCAGAGCTGATAATCAGGTCAAGATTCGGG |

| | |
|---|---|
| | GTTTTCGTGTTGAATTGGGTGAAGTAGAAACTGCCCTTACCAATTGTCCCCCAGTGTCAGATGCCCTCGCTATGGTTCGTGAGGATCGACCGGGCGAAAAATTCCTAG<br>TGGCCTATGTGGTTGGTCAGGATTCTATGGATACTGATGCTCTGCGTGCCCAGCTCGTTAATGATTTACCTCCATACTTGATC |
| Ltx-F5 | CTCGTTAATGATTTACCTCCATACTTGATCCCTGGCGCTATTGTCACTTTGAAGAAATTTCCCCTGACAGTGAACGGTAAAATAGATCGTAAGGCTCTCCCAGTTCCAAA<br>GTTCACTTCTATTGCTGAAGGGCGTGTTCCCAAAACTCCTACCCAGATTGGTTTATGTACTCTGTTTGCAGAACTGTTGGGTGTTGATCAAGTCACGATCGACGATCCG<br>TTTTTTGCCCTTGGAGGTCACTCGTTGCTAGCCACACAATTAGTGAGTCAGATTCGGCAGCGTTTTGGGATAAGTTTGCCAGTGCAAACTGTATTCGAGCGACAGACG<br>GTGGCTGACCTGGCAATAGTAGTAGATCGGGAAGCACCTATAGTTTCTACTTCCATTGACCTTGCTGCTGAAGTTGTTCTTAATCCGCAGATTGCTCCCTACCAGAGTC<br>GTCCAGTTGAACTCGATAGGAATACTCACCCAGCCTCGGTGTTATTAACCGGAGCAACTGGCTTTTTGGGGGCATATTTGCTTTATGAACTTCTGAAGCAGACTGATGC<br>TAATGTATTTTGCTTGGTGCGCTCCAACCACTCTGAAGCTGCCTATCAACGCATCCATTCGACTCTCAAATTTTATCAATTATGGTCTGAATCTTGGAGATCGCGAATCAT<br>TCCAGTGTGTGGCGATCTCTCCCAGCCAAGTTTAGGCTTATCCGCTGAGGAATTCAGCAAGCTCACAGAATTAATTGATGCTATCTATCACAATGGAGCCCAGGTCAGT<br>GCTATTGAACCCTATACATACCTTAAACCAACAAATGTATTGGGGACATCAGAGTTACTGGACTTTGCTGCGCGATGCCGCGTGAAGCCATTGCACTTTGTTTCAACAG<br>CTGCAGTGGCTGTAAGTTCTAAAGGTAATCCTGATATAATTTATGAGAATTTTAGACTTGGAGCAGATTCTGTATTACCCAGTGGATATGTCTCTAGTAAATGGGTCGCTG<br>AAGAGTTAGTGTGGGTGGCATCAGACCGTGGCCTCCCCGTTACTGTCCACCGCCCAGGCCGAATTAGTGGTGATACCACTACTGGTATTGGGAATACTGATGATACATT<br>CTGGCAAATAGTCAGGGCAATAGTTGTTCTAGGGGTTGTCCCAGATATTGTTTATCAAGATGATGCCGGGATCGATTTAATGCCAGTCGATCGCGTTGCAAGTGCAATT<br>GTCCATCTATCCCGTCATAAACAATCCATAAGCAAGGTGCATCACCTCACCTGTCCTACAATAGTCAAGCTAGATGTAGTGTTTAATGAACTCTCTAAATTGGGCTACCA<br>GTTAACCACGGTATCCTACTCTGAATGGGTTAAGCAATTAGAACAGTACGTCGATCAAGCCCCAGGGGGGCACTCCCTAGCATCAGCTACCGTCCTTAGTCGTACATTA<br>CCCAAGCTCATAGAACTCAGTCAAATCTGCTTCGACCAAAGCAACACCTTAACGGGTTTAGCTGAAGCACCATTTAAGTTTCCTTGTATTGATCGTCATCTTGTCCGTG<br>GATACCTCACTTATTTCATCAACTCTAAATTCTTCCCTCAAATAACTTATACAGGAAAGTAACAGCTTTCTTGTACAAAGTTGGCATTATAA |

**Table S2**: Synthetic gene fragments to reconstruct *ltxA in vivo*

| Primer name | Sequence | Description |
|---|---|---|
| ES-LtxA-F | ATGATTATGAATCAACCTTGGA GTG | Forward primer to amplify the *ltxA* gene that is used for identification and sequencing purposes |
| ES-LtxA-M-R | GTCAATTCAAACTGACCCGGAT TC | Reverse primer that binds at the 3303 bp of *ltxA*. This was used for the identification of RNA truncation. |
| ES-LtxA-M-F | GTTCTATTGAATCCGGGTCAGT TTGAATTG | Forward primer that binds at the 3303 bp of *ltxA*. This was used for the identification of RNA truncation. |
| SJ-LtxA-M2-R | CAAATCAGTACGATACTCAACC | Reverse primer that binds at the 4290 bp of *ltxA*. This was used for the identification of RNA truncation. |
| ES-LtxA-R | TTACTTTCCTGTATAAGTTATTT GAGGGA | Reverse primer to amplify the *ltxA* gene that is used for identification and sequencing purposes |
| ES_SJ_attR1-LtxB-F | CAT ATC CAG TCA TAT TGG CAT GAC AAA TCC TTT TGC AG | Forward primer to amplify the *ltxB* gene with overhangs that match with the *Not*I restriction site. |
| ES_SJ_camR-LtxB-R | CTG GGG TGC CTA ATG CTT ACC ACT CAG CAG GTA AC | Reverse primer to amplify the *ltxB* gene with overhangs that match with the *Not*I restriction site. |
| ES_SJ_attR1-LtxC-F | CAT ATC CAG TCA TAT TGG CAT GAA TTC AAA GAT CGC TG | Forward primer to amplify the *ltxC* gene with overhangs that match with the *Not*I restriction site. |
| ES_SJ_camR-LtxC-R | AAG CCT GGG GTG CCT AAT GCT TAT GAC CGT TTA TAG ACT C | Reverse primer to amplify the *ltxC* gene with overhangs that match with the *Not*I restriction site. |
| SJ-sfp-F | ATGAAGATCTACGGAATATACA TGGATCG | Forward primer to amplify the *sfp* gene that is used for identification and sequencing purposes |
| SJ-sfp-R | CTACAGCAACTCCTCGTAAGAA ACC | Reverse primer to amplify the *sfp* gene that is used for identification and sequencing purposes |
| SJ-hetI-F | ATGTTGCAGCACACATGGC | Forward primer to amplify the *hetI* gene that is used for identification and sequencing purposes |
| SJ-hetI-R | CAGCCGAAGTTTTGGCACTAT | Reverse primer to amplify the *hetI* gene that is used for identification and sequencing purposes |
| ES-pamyB-F | TATATGGCGGGTGGTGGGC | Forward primer to amplify P*amyB* promoter onwards that is used for identification and sequencing purposes |

| | | |
|---|---|---|
| ES-tamyB-R | TTCCGTTCCTTTGCTTTCTGCCGAGC | Reverse primer to amplify P*amyB* terminator that is used for identification and sequencing purposes |
| pet22-ltxB-F | TTTTGTTTAACTTTAAGAAGGAGATATAATGACAAATCCTTTTGCAGAC | Forward primer for *ltxB* with overhangs to RBS of pET22b(+) vector |
| pet22-ltxB-R | AAAGTTACCTGCTGAGTGGCACCACCACCACCACCACTGAGATCCGGC | Reverse primer for *ltxB* with overhangs to his tag of pET22b(+) vector |
| pet22-F | CACCACCACCACCACCACT | Forward primer for his tag of pET22b(+) vector |
| pet22-R | TATATCTCCTTCTTAAAGTTAAACAAAATTATTTCTAGAGGGG | Reverse primer for RBS of pET22b(+) vector |
| pET15b-ltxB-F | CAGCGGCCTGGTGCCGCGCGGCAGCCATATGATGACAAATCCTTTTGCAG | Forward primer to amplify *ltxB* with 30 bp overhang to pET15b+ vector |
| pET15b-R | CATATGGCTGCCGCGC | Forward primer to amplify the pET15b+ vector |
| pET15b-ltxB-R | TTCCTTTCGGGCTTTGTTAGCAGCCGGATCTTACCACTCAGCAGGTAAC | Reverse primer to amplify ltxB with 30 bp overhang to pET15b+ vector |
| pET15b-F | GATCCGGCTGCTAACAAAGC | Reverse primer to amplify pET15b+ vector |
| pre-pamyB-F | CATGGTGTTTTGATCATTTTA | Forward primer anneals to a sequence upstream of the P*amyB* promoter in the pTYGxxx vector |
| post-tamyB-R | GATAACAATTTCACACAGG | Reverse primer anneals to a sequence downstream of the T*amyB* terminator in the pTYGxxx vector |

**Table S3**: Oligonucleotides used in this study

#### 1.1 Plasmids used in this study

Plasmids used in this study are listed in Table S4 and the corresponding plasmid maps are shown in Figure S1. Briefly, *ltxA* was constructed from five overlapping fragments of synthetic DNA (Table S2) into pE-YA using yeast homologous recombination; subsequently, Gateway™ cloning was used to clone *ltxA* into pTYGSarg fungal expression vector under the *amyB* promoter. *ltxB* and *ltxC* were individually cloned into expression vectors pTYGSniaD and pTYGSmet under the *amyB* promoter, using the Takara In-Fusion® Snap Assembly master mix, following the manufacturer's instructions. *Sfp* and *hetI* genes were ordered as synthetic genes cloned in the pTwist-ENTR vector, and were subsequently transferred to pTYGSade using LR recombination™ cloning.

| **Plasmid name** | **Description** | **Reference** |
|---|---|---|
| pE-YA | *E. coli*/yeast cloning plasmid (*kan*R, pUC ori, 2μ ori, *URA3*) | Lazarus et al., 2014 |
| pTYGSarg | *E. coli*/yeast/*A. oryzae* shuttle vector and expression plasmid (P*amyB*, P*adh*, P*eno*, P*gdpA*, *amp*R, ColE1, 2μ ori, *URA3*, *ccdB*, *argB*) | Pahirulzaman et al., 2012 |
| pTYGSade | *E. coli*/yeast/*A. oryzae* shuttle vector and expression plasmid (P*amyB*, P*adh*, P*eno*, P*gdpA*, *amp*R, ColE1, 2μ ori, *URA3*, *ccdB*, *adeA*) | |
| pTYGSmet | *E. coli*/yeast/*A. oryzae* shuttle vector and expression plasmid (P*amyB*, P*adh*, P*eno*, P*gdpA*, *amp*R, ColE1, 2μ ori, *URA3*, *ccdB*, *sC*) | |
| pTYGSniaD | *E. coli*/yeast/*A. oryzae* shuttle vector and expression plasmid (P*amyB*, P*adh*, P*eno*, P*gdpA*, *amp*R, ColE1, 2μ ori, *URA3*, *ccdB*, *niaD*) | |
| pE-YA+LtxA | *E. coli*/yeast cloning plasmid (*kan*R, pUC ori, 2μ ori, *URA3*, *ltxA*) | This study |
| pTYGSarg+LtxA | *E. coli*/yeast/*A. oryzae* shuttle vector and expression plasmid (P*amyB*, P*adh*, P*eno*, P*gdpA*, *amp*R, ColE1, 2μ ori, *URA3*, *argB*, *ltxA*) | This study |
| pTYGSniaD+LtxB | *E. coli*/yeast/*A. oryzae* shuttle vector and expression plasmid (P*amyB*, P*adh*, P*eno*, P*gdpA*, *amp*R, ColE1, 2μ ori, *URA3*, *niaD*, *ltxB*) | This study |
| pTYGSmet+LtxC | *E. coli*/yeast/*A. oryzae* shuttle vector and expression plasmid (P*amyB*, P*adh*, P*eno*, P*gdpA*, *amp*R, ColE1, 2μ ori, *URA3*, *sC*, *ltxC*) | This study |
| pTYGSade+HetI | *E. coli*/yeast/*A. oryzae* shuttle vector and expression plasmid (P*amyB*, P*adh*, P*eno*, P*gdpA*, *amp*R, ColE1, 2μ ori, *URA3*, *adeA*, *hetI*) | This study |
| pTYGSade+sfp | *E. coli*/yeast/*A. oryzae* shuttle vector and expression plasmid (P*amyB*, P*adh*, P*eno*, P*gdpA*, *amp*R, ColE1, 2μ ori, *URA3*, *ccdB*, *adeA*, *sfp*) | This study |
| pET-22b(+)-ltxB | *E. coli* protein expression plasmid (T7 promoter, *lac*, T7 gene, *pelB*, C-terminal 6xHis tag, T7 terminator, *amp*R/*bla*, ColE1/pMB1/pBR322/pUC origin, f1 origin, *lacI*, *rop*, *bom*, *ltxB*) | This study |
| pET-15b-ltxB | *Modified* pET-15b with spectinomycin-resistant gene, *E. coli* protein expression plasmid (T7 promoter, *lac*, T7 gene, *pelB*, N-terminal 6xHis tag, T7 terminator, *aadA*, ColE1/pBR322/pUC origin, f1 origin, *lacI*, *rop*, *bom*, *ltxB*) | This study |

| | | |
|---|---|---|
| pTYGSarg+LtxC-CO-eGFP | *E. coli*/yeast/*A. oryzae* shuttle vector and expression plasmid (P*amyB*, P*adh*, P*eno*, P*gdpA*, *amp*R, ColE1, 2μ ori, *URA3*, *argB*, *ltxC*-CO, *eGFP*) | This study |

**Table S4**: Plasmids used in this study.

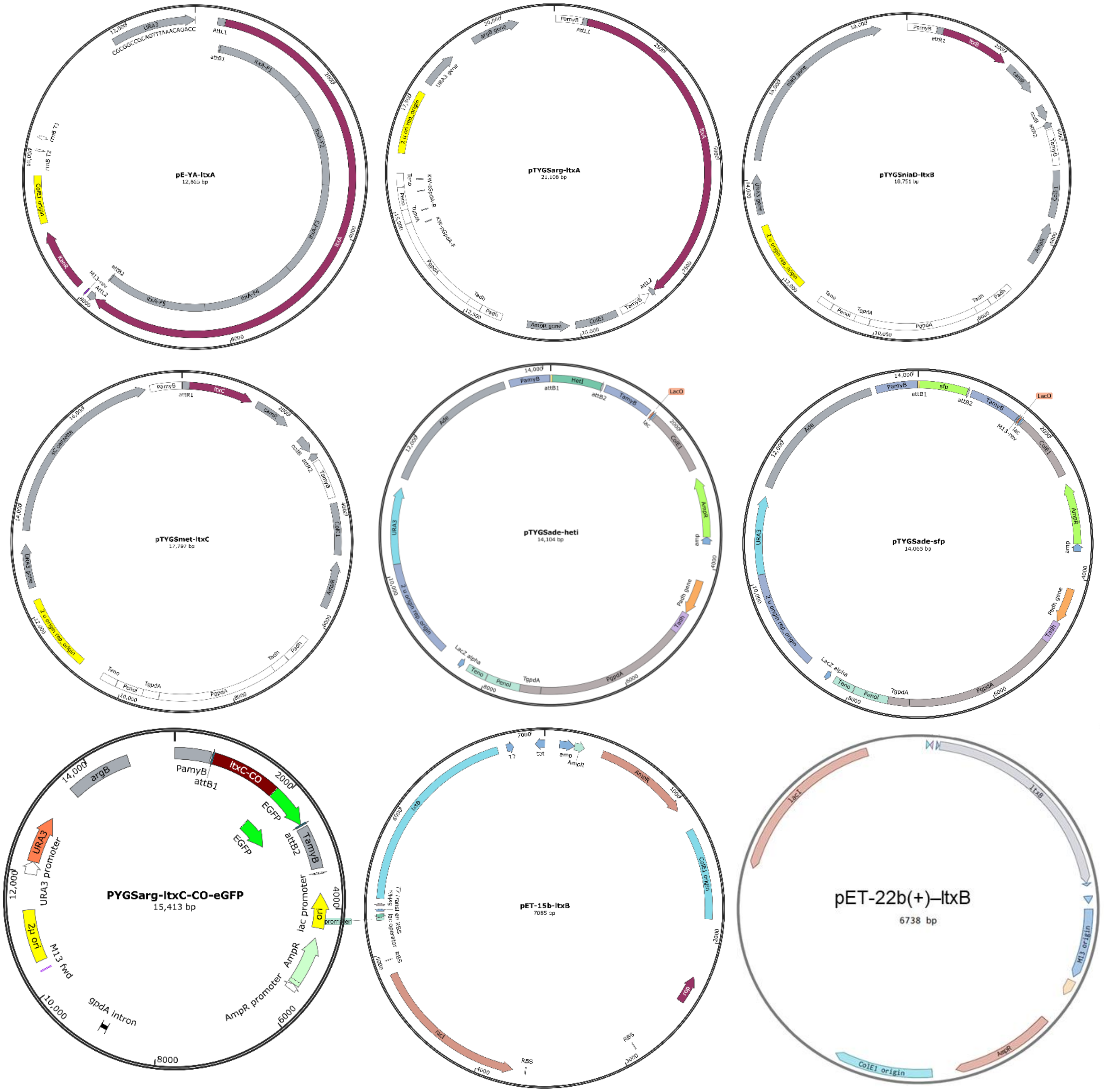

**Figure S1**: Maps of plasmids used in this study.

### 1.2 Strains, media, and solutions used in this study

Microbial strains used in this study are listed in Table S5. All media were prepared in deionized water and autoclaved at 121 °C for 20 min (Table S6).

| Strain name | Genotype / Description |
|---|---|
| *A. oryzae* NSAR1 | *argB-, adeA-, sC-, niaD-* |
| *E. coli* TOP10 | F⁻ *mcrA*, Δ(*mrr-hsdRMS-mcrBC*), *φ80lacZ*ΔM15, Δ*lacX74*, *recA1*, *araD139*, Δ(*ara-leu*)7697, *galU*, *galK*, *rpsL* (Str^R), *endA1*, *nupG* |
| *E. coli ccdB* survival | F- mcrA Δ(mrr-hsdRMS-mcrBC) Φ80lacZΔM15 ΔlacX74 recA1 araΔ139 Δ(ara-leu)7697 galU galK rpsL (StrR) endA1 nupG fhuA::IS2 |
| DE3 *E. coli* | F⁻ *ompT*, *gal*, *dcm*, *lon*, *hsdS*B(rB– mB–) λ(DE3 [*lacI*, *lacUV5-T7*p07, *ind1*, *sam7*, *nin5*]), [*malB*$^{+}$]K-12(λ$^{S}$) |
| *S. cerevisea* INScv | *MATa his3D1 leu2 trp1-289 ura3-52 MAT his3D1 leu2 trp1-289 ura3-52* |
| *A. oryzae* + *ltxABC+sfp* | *A. oryzae* NSAR1 expressing *ltxA*, *ltxB*, *ltxC*, and *sfp*. |
| *A. oryzae ltxABC+hetI* | *A. oryzae* NSAR1 expressing *ltxA*, *ltxB*, *ltxC*, and *hetI*. |
| *A. oryzae* + *ltxC*-co | *A. oryzae* NSAR1 expressing codon optimized *ltxC* |
| DE3 *E. coli* + *ltxB*(C) | DE3 *E. coli* expressing *ltxB* with a C-terminal 6xHis-tag |
| DE3 *E. coli* + *ltxB*(N) | DE3 *E. coli* expressing *ltxB* with an N-terminal 6xHis-tag |

**Table S5**: Strains used in this study

| Medium Name | Ingredients |
|---|---|
| CZD/S agar | 35 g/L Czapek Dox broth, 182.2 g/L D-sorbitol, 1 g/L ammonium sulphate, 0.5 g/L adenine, 1.5 g/L-methionine, 15 g/L agar |
| CZD/S soft agar | 35 g/L Czapek Dox broth, 182.2 g/L D-sorbitol, 1 g/L ammonium sulphate, 0.5 g/L adenine, 1.5 g/L-methionine, 8 g/L agar |
| CZD/S1 agar | 35 g/L Czapek Dox broth, 182.2 g/L D-sorbitol, 1 g/L ammonium sulphate, 1.5 g/L-methionine, 15 g/L agar |
| CZD/S1 soft agar | 35 g/L Czapek Dox broth, 182.2 g/L D-sorbitol, 1 g/L ammonium sulphate, 1.5 g/L-methionine, 8 g/L agar |
| CZD/S1 agar w/o methionine | 35 g/L Czapek Dox broth, 182.2 g/L D-sorbitol, 1 g/L ammonium sulphate, 15 g/L agar |
| CZD/S1 soft agar w/o methionine | 35 g/L Czapek Dox broth, 182.2 g/L D-sorbitol, 1 g/L ammonium sulphate, 8 g/L agar |
| DPY medium | 20 g/L dextrin from potato starch, 10 g/L polypeptone, 5 g/L $KH_2PO_4$, 0.5 g/L $MgSO_4 \cdot H_2O$ |
| GN medium | 20 g/L D (+)-glucose monohydrate, 10 g/L nutrient broth No. 2 from oxoid (Thermo Scientific) |
| LB agar | 5 g/L yeast extract, 10 g/L tryptone, 5 g/L NaCl, 15 g/L agar |

| | |
|---|---|
| SM-ura agar | 1.7 g/L yeast nitrogen base, 20 g/L D (+)-glucose monohydrate, 5 g/L ammonium sulphate, 0.77 g/L complete supplement mixture minus uracil (Q biogene), 25 g/L agar |
| SOC medium | 937.5 mL/L SOB medium, 12.5 mL/L $MgCl_2$, 50 mL/L glucose (20 %) components were autoclaved separately and sterile filtrated after mixing |
| YPAD agar | 10 g/L yeast extract, 10 g/L tryptone, 0.3 g/L adenine, 20 g/L D (+)-glucose monohydrate, 15 g/L agar |
| YPAD medium | 10 g/L yeast extract, 10 g/L tryptone, 0.3 g/L adenine, 20 g/L D (+)-glucose monohydrate |
| Czapek Dox Broth | 3.50% (w/v) Czapek Dox broth (Difco) was added to 100 mL of Mili-Q water, dispensed into a 200 mL media bottle and autoclaved. |
| Starch medium | 20.0 g/L potato starch using repeated microwave bursts, 10.0 g/L polypeptone, 5.00% (v/v) Solution A (4.00% sodium nitrate, 4.00% potassium chloride, 0.93% magnesium sulfate hexahydrate, 0.02% iron(II) sulfate heptahydrate) and 5.00% (v/v) Solution B (2.00% dipotassium phosphate). |
| Glucose medium | 20 g/L D(+)-glucose monohydrate and 10.0 g/L polypeptone, 5.00% (v/v) Solution A (4.00% sodium nitrate, 4.00% potassium chloride, 0.93% magnesium sulfate hexahydrate, 0.02% iron(II) sulfate heptahydrate) and 5.00% (v/v) Solution B (2.00% dipotassium phosphate). |

**Table S6**: Recipes of media used in this study.

| **Solution** | **Ingredients** |
|---|---|
| *E. coli* lysis buffer | 50mM Tris pH 8.0, 100mM NaCl, 1% Triton X-100, in 100 mL of Milli-Q water |
| 1.5 M Tris-HCl, pH 8.8 | Diluted Tris-HCl 2M solution, Adjust pH to 8.8 |
| 0.5 M Tris-HCl, pH 6.8 | Diluted Tris-HCl 2M solution, Adjust pH to 6.8 |
| 30% acrylamide/bisacrylamide solution | 29:1 g acrylamide: bisacrylamide in 100 mL of Milli-Q water |
| 10% SDS | 10g of SDS solid in 100mL of Milli-Q water |
| 10% ammonium persulfate (APS) | 100mg of APS solid in 1 mL of Milli-Q water. The solution was made fresh before use. |
| Running buffer | Diluted 10X stock solution (Tris-Glycine, Biorad) in Milli-Q water |
| Coomassie stain | 2 g Coomassie Blue R-250 in 250 mL water, 75 mL of glacial acetic acid, and 500 mL ethanol, in 1 L of Milli-Q water |
| Destain solution | 50% methanol, 10% acetic acid, 40% milli-Q water |
| Wash buffer-1 | 50 mM Tris-HCl (pH 8.0), 500 mM NaCl, 1% Triton X-100, and 10 mM imidazole. |
| Wash buffer-2 | 50 mM Tris-HCl (8.0), 500 mM NaCl, 0.03% DDM, and 25 mM imidazole. |
| Elution buffer | 50 mM Tris-HCl (pH 8.0), 500 mM NaCl, 0.03% DDM, and 250 mM imidazole. |

| | |
|---|---|
| 10x TBS buffer | 10x TBS buffer, 24.2g of Tris base, and 88 g of NaCl were dissolved in 1 L of Milli-Q water, and the pH was adjusted to 7.6 with HCl. |
| Protein transfer buffer to PVDF membrane | 303 mg of Tris base, 1.5 g of glycine, 20 mL of methanol, and 80 mL of MiliQ. pH adjusted to be between 8.5 and 9.0 |
| TBS/T buffer | 10x TBS buffer was diluted to 1x TBS buffer with Mili-Q water and add 0.1% Tween-20. |
| Blocking buffer for Western blot | 5 g nonfat dry milk in 100 mL of 1x TBS buffer with 0.1% Tween-20 (TBS/T). |

**Table S7**: Recipes of buffers and other chemicals used in this study.

#### 1.3 Fungal cultivation and extraction methods

A spore suspension of *A. oryzae* NSAR1 / transformants were grown on DPY agar plates or inoculated into 100 mL DPY liquid medium in a 500 mL baffled flask and incubated for at 28 °C with 110 rpm shaking. After 7 days, the liquid supernatant was extracted twice with an equal volume of ethyl acetate. Combined organic layers were dried over $MgSO_4$ and the solvent was removed under reduced pressure. The organic residue was dissolved in LCMS grade methanol to a concentration of 10 mg/mL, filtered over glass wool and analyzed by LCMS.

**Timecourse experiment**

*A. oryzae* transformant-18 was grown in Czapek dox broth, glucose broth, Czapek dox broth-starch, glucose -starch and DPY media. From day 4 to 11 days, growth media and cells were extracted using ethyl acetate. The extracted metabolites were combined based on growth media and analyzed using an LC-TripleTOF® 6600+ mass spectrometer. In the analysis, expected *m/z* values were evaluated using exact mass, previously reported m/z values, and m/z values obtained from running standards in this LC-MS/MS method.

**Feeding/biotransformation of commercial substrates**

Commercial ILV and LTXA (Cayman Chemical) substrates were dissolved in 100% ethanol to make 0.5 mg/mL LTXA and 2 mg/mL ILV stock solutions respectively. 50 µL of LTXA stock solution was added to 500 mL Erlenmeyer flasks containing *A. oryzae* NSAR1 that had been cultivated for 1 day. After 7 days, the cells and media were extracted separately as described above and analyzed by LCMS. 50 µL of ILV stock solution was added to 500 mL Erlenmeyer flasks containing *A. oryzae* NSAR1, *A. oryzae-ltxABC+hetI* (ID:18), and *A. oryzae-ltxABCD+hetI* (ID:21) strains separately that had been cultivated for 1 day. After 7 days, the cells and media were extracted separately, as described above, and analyzed by LC-MS.

### 1.4 Transformation methods

#### Chemically competent *E. coli* cells

Chemically competent *E. coli* cells were thawed on ice and gently mixed with 1–10 ng of plasmid DNA in a sterile microcentrifuge tube. Transformation proceeded following the manufacturer's instructions. The transformation mixture was shaken at 37°C (220 RPM) for one hour, and approximately 50 µL was plated onto LB agar with added selective antibiotic and incubated overnight.

#### Electrocompetent *E. coli* cells

Thawed electrocompetent cells were combined with DNA (1-100 ng), loaded in cold electroporation cuvettes, and given a 1.8 kV pulse according to the manufacturer's protocol. Following shock, 950mL of LB media was added. The transformation mixture was shaken at 37°C (220 RPM) for one hour, and approximately 50 µL was plated onto LB agar with added selective antibiotic and incubated overnight.

#### Yeast

*S. cerevisiae* was streaked out on YPAD agar and incubated at 30 °C for 3-5 days. A single colony was transferred into 10 mL YPAD medium and incubated overnight at 30 °C with 200 rpm shaking. This starter culture was added to 40 mL of YPAD medium in a 250 mL flask and incubated for an additional 4.5 hr at 30 °C with shaking at 200 rpm. The culture was harvested by centrifugation at 4600 RPM for 5 min. The pellet was washed with 25 mL $ddH_2O$ and centrifugation was repeated. The pellet was resuspended in 1 mL of 100 mM LiOAc and centrifuged at 14000 RPM for 15 seconds. The supernatant was removed, and the pellet was resuspended in 500 µL of 100 mM LiOAc. The cell suspension was aliquoted to 50 µL and transferred into separate 1.5 mL reaction tubes. For each sample, one aliquot was centrifuged at 14000 RPM for 15 s and the supernatant was removed. The pellet was dissolved in the transformation mixture consisting of 240 µL PEG solution (50 % (w/v) polyethylene glycol 3350),36 µL LiOAc (1 M), 50 µL denatured salmon sperm DNA (2 mg/mL in TE buffer), 34 µL DNA master mix containing the linearized vector and desired inserts obtained by PCR in equimolar concentration. The synthetic oligos contain 30 bp overlaps at both 5' and 3' with the cut sites of the vector fragments to facilitate homologous recombination. Cells were first incubated for 30 min at 30 °C, then for 40 min at 42 °C. Cells were pelleted by centrifugation at 6000 RPM for 15 s, and the supernatant was removed. The pellet was resuspended in 1 mL $ddH_2O$ and 250 µL was spread on selective SM-Ura plates, which were incubated for four days at 30 °C.

Constructed plasmid DNA was extracted from yeast cells using a Zymoprep™ Yeast Plasmid Miniprep II kit (Zymo Research, Orange, California, USA) and transformed into *E. coli ccdB* Survival cells by the standard heat shock method for amplification.

***Aspergillus oryzae***

A spore suspension collected from a 7-day-old *A. oryzae* NSAR1 DPY plate was used to inoculate 50 mL (250 mL flask) of GN liquid culture and incubated for 16 h (28 °C, 110 rpm). Cells were collected by filtration over sterile Miracloth, washed with 0.8M NaCl, and suspended in 10 mL of filter-sterilized water. *A. oryzae* NSAR1 protoplasting solution (10 mg/mL lysing enzyme from *Trichoderma harzianum* or Yatalase, 0.8M NaCl, 10 mM $CaCl_2$). The suspension was incubated for 4 hrs at ambient temperature with gentle shaking. Protoplasts were released by pipetting, collected by centrifugation (3000 × g, 5 min) and directly suspended in the required amount of fungal transformation solution I (10mM CaCl2, 0.8M NaCl and 50mM Tris-HCl at pH 7.5). Vector DNA (≥1 µg in 10 µL of ddH2O) was mixed with 100 µL protoplasts and incubated on ice for 5 min. One millilitre of fungal transformation solution II (10mM CaCl2, 0.8M NaCl and 50 mM Tris-HCl at pH 7.5, 60% (w/v) PEG3350) was added and the mixture was incubated at ambient temperature for 20 min. 5 mLs of molten selective soft agar (CZD/S, CZD/S1 or CZD/S1 w/o methionine) was added and the mixture was poured over selective agar plates (CZD/S, CZD/S1 or CZD/S1 w/o methionine). Plates were incubated at 28 °C until colonies appeared, which were transferred to secondary plates of the respective selective agar. Vigorously growing colonies were transferred onto a third plate selective plate.

### 1.5 gDNA / RNA extraction and PCR methods

*A. oryzae* colonies were screened for gene insertion using the Platinum direct PCR kit (Invitrogen) following the manufacturer's instructions and gene specific primers (Table S3). Prior to starting RNA extraction, all apparatus and surfaces were treated with RNase AWAY, wiped with a paper towel, and let it dry for a minute. Transformants were grown in DPY broth for 3 days, and the cells were separated using gravity filtration on sterilized filter paper. Cells were immediately frozen with liquid nitrogen in sterilized mortars and then directly ground with a pestle. Approximately 80 mg of the sample was placed into frozen microcentrifuge tubes. RNA extraction was performed using the Qiagen RNeasy Mini Kit, following the manufacturer's instructions. The homogenate was then transferred to a QIAshredder spin column (Qiagen), again following the manufacturer's instructions.

Following DNA digestion, the RNA was purified using the Monarch RNA Cleanup Kit (New England Biolabs). For cDNA synthesis, 1000 ng of purified RNA was used as input in a 20 µl reverse transcription reaction using the LunaScript RT SuperMix Kit (New England Biolabs). The resulting cDNA was either used immediately for downstream applications or stored at –20°C until use.

**PCR**

PCR was carried out using NEB Q5 master mix or Takara PrimeStar master mix. Previously prepared cDNA 2 μL was used as a template for a 50 μL PCR reaction, and agarose gel (0.8%) electrophoresis was carried out to visualize the DNA bands.

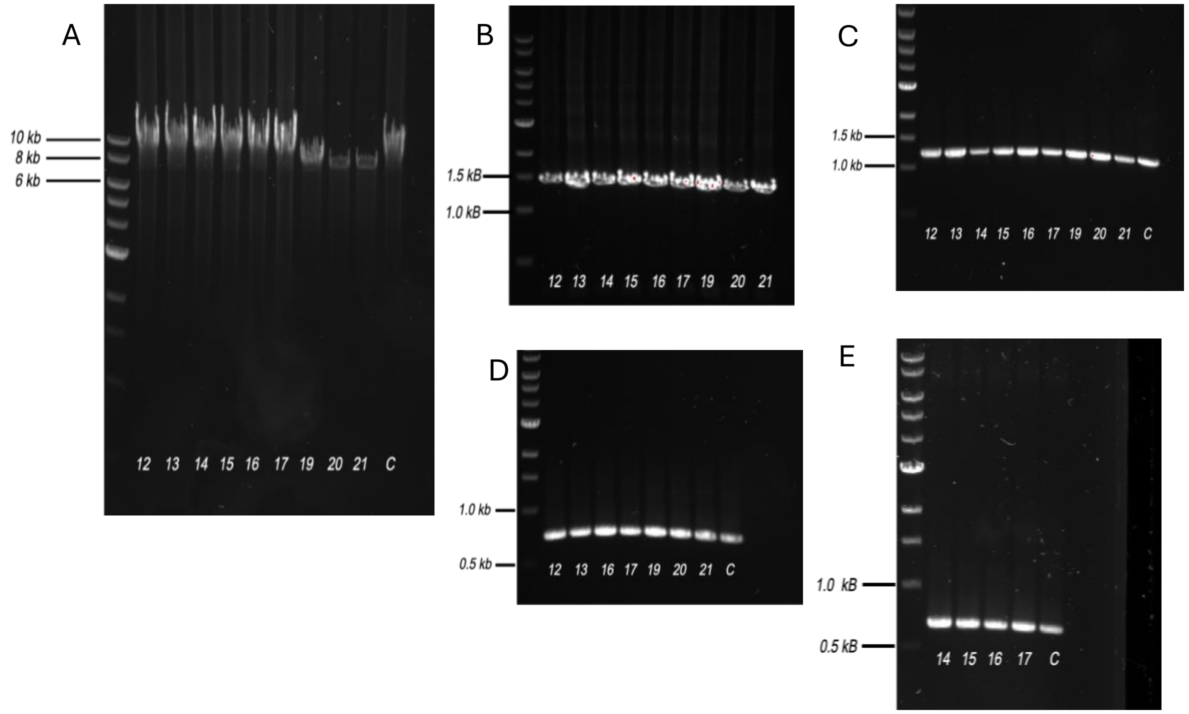

**Figure S2**: PCR screening of *A. oryzae* transformants integrating: A) *ltxA*; B) *ltxB*; C) *ltxC*; D) *hetI*; and E) *sfp*. Lanes are labeled according to transformant ID; C = plasmid template used as positive control

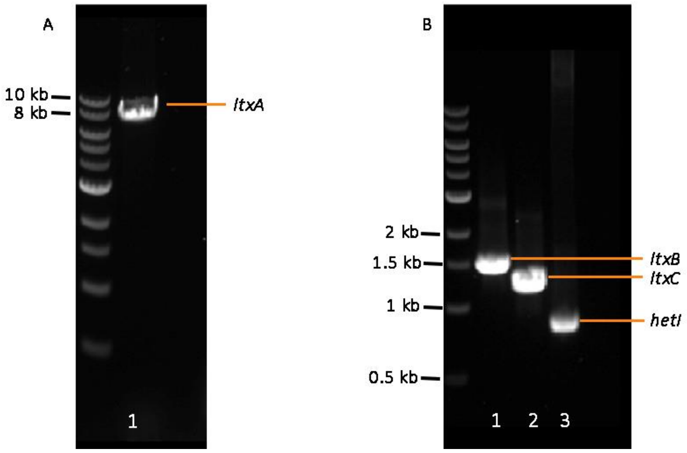

**Figure S3**: *A. oryzae* transformant 18 was confirmed as containing *ltxA*-*C* and *hetI* genes and was subsequently used for RT-PCR, timecourse and media studies.

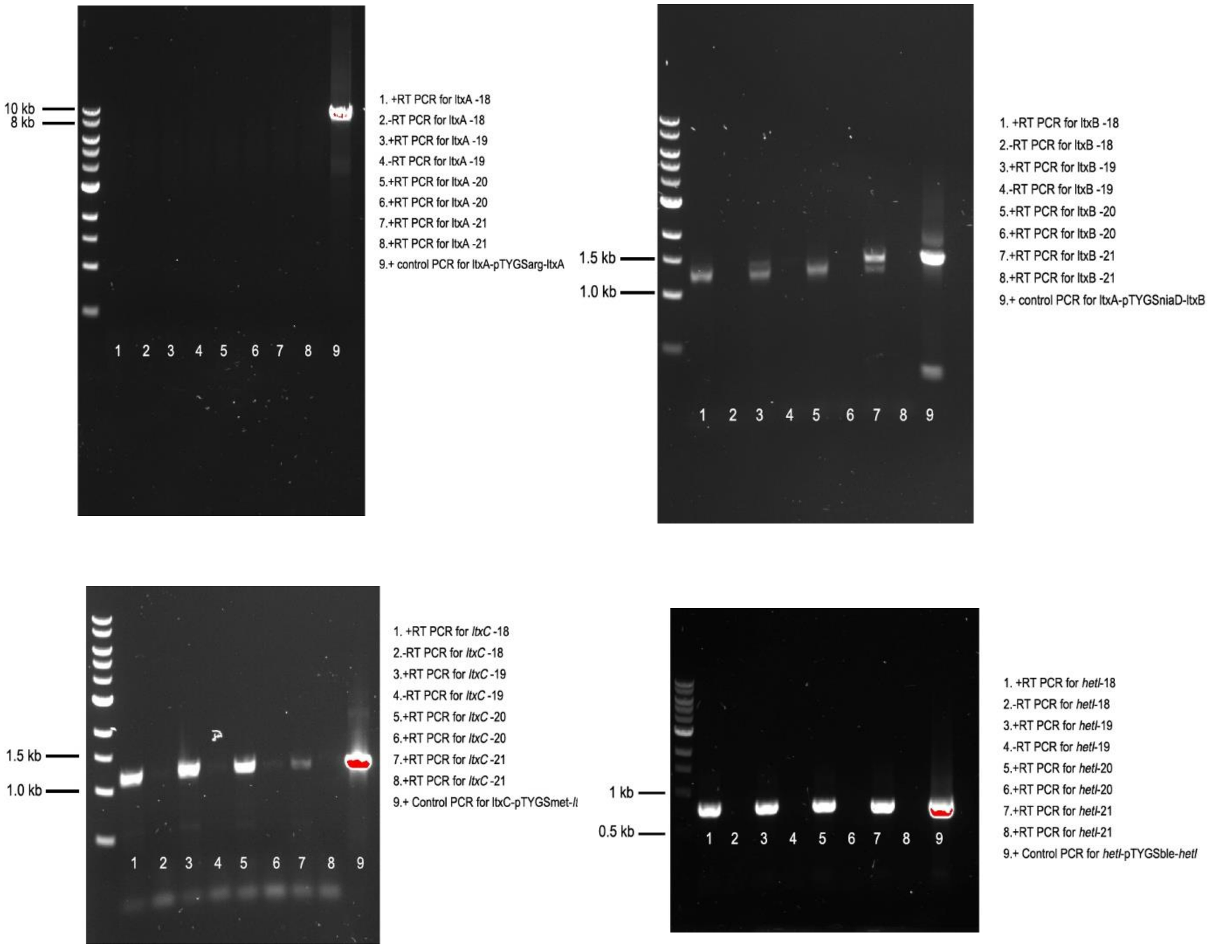

**Figure S4**: Transcription analysis of *ltxA-C* and *hetI* in transformants 18-21.

To examine RNA truncation, the cDNA from transformant 18 was subjected to PCR amplification. This experiment was carried out using multiple reverse primers that anneal to the *ltxA* cDNA (Figure S5 ). Each reverse primer was used in combination with the ltxA-F primer, which anneals at the beginning of ltxA, and RNA truncation was observed between 3304 bp and 4290 bp (Figure S5). All transcripts were sequenced and verified by Plasmidsaurus, and no mutations were detected.

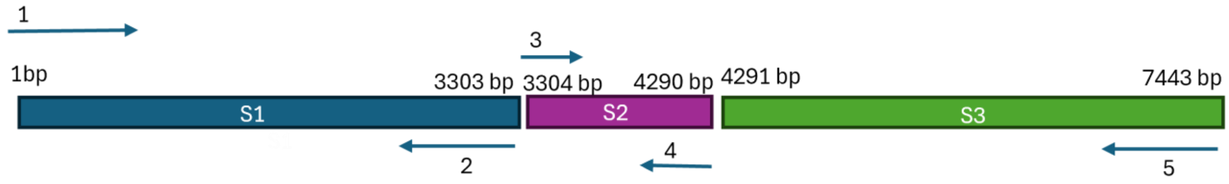

**Figure S5**: PCR strategy to determine where *ltxA* is truncated during transcription.

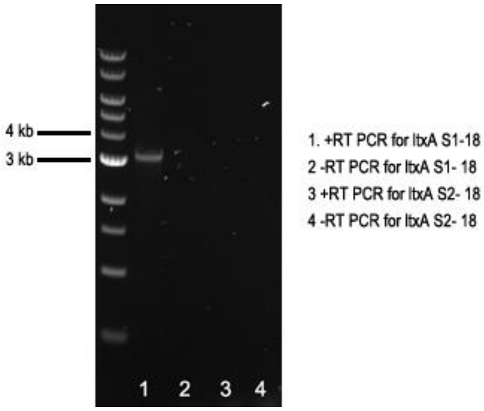

`

**Figure S6**: *ltxA* $1^{st}$ first segment (S1: 1 bp-3303 bp) had positive transcription. *ltxA* $2^{nd}$ segments (S2: 3303bp-4290 bp) was not detected.

#### 1.6 Bioinformatics

Enzyme localization predictions in fungal hosts were performed using PSORTII and WoLF PSORT (Table S8). Potential intron sequences/splice sites were searched for using FGENESH (Figures S7 - 9). Sequence alignments were performed using Geneious™ and proteins were modelled using AlphaFold server 3.0 (Abraham et al., 2024).

| **Enzyme Name** | **PSORTII localization prediction** (Nakai et al., 1999) | **WoLF PSORT localization prediction** (Horton et al., 2007) | **TargetP 2.0** (Armenteros et al., 2019) |
|---|---|---|---|
| LtxA | 44% ER | 16 plasma membrane | Other (no targeting peptide) |
| LtxB | 30% nuclear | 9 nuclear | Other |
| LtxC | 35% nuclear; 30% mitochondrial | 8 cytoplasm / nuclear | Other |

**Table S8:** *In silico* predictions of LTX biosynthetic enzyme localization in a fungal host

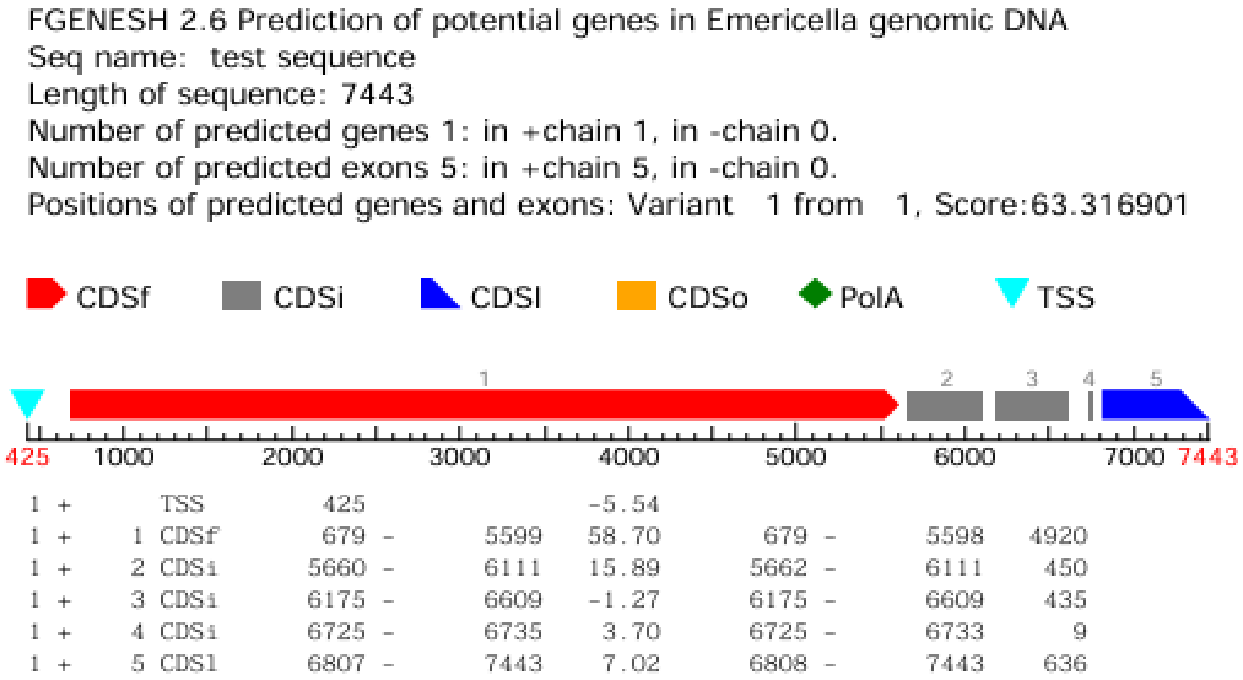

**Figure S7**: Exon and intron predictions of *ltxA* processed by *Emericella* (*Aspergillus*) sp. using FGENESH.

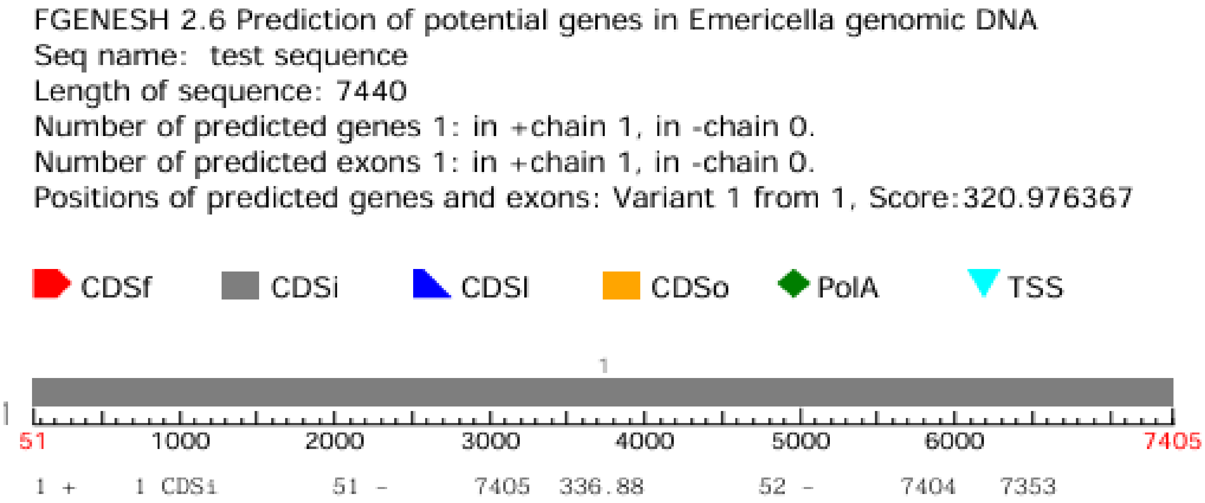

**Figure S8**: Exon and intron predictions of codon optimized *ltxA* processed by *Emericella* (*Aspergillus*) sp. by FGENESH. No intron sequences are predicted however the predicted protein sequence is truncated.

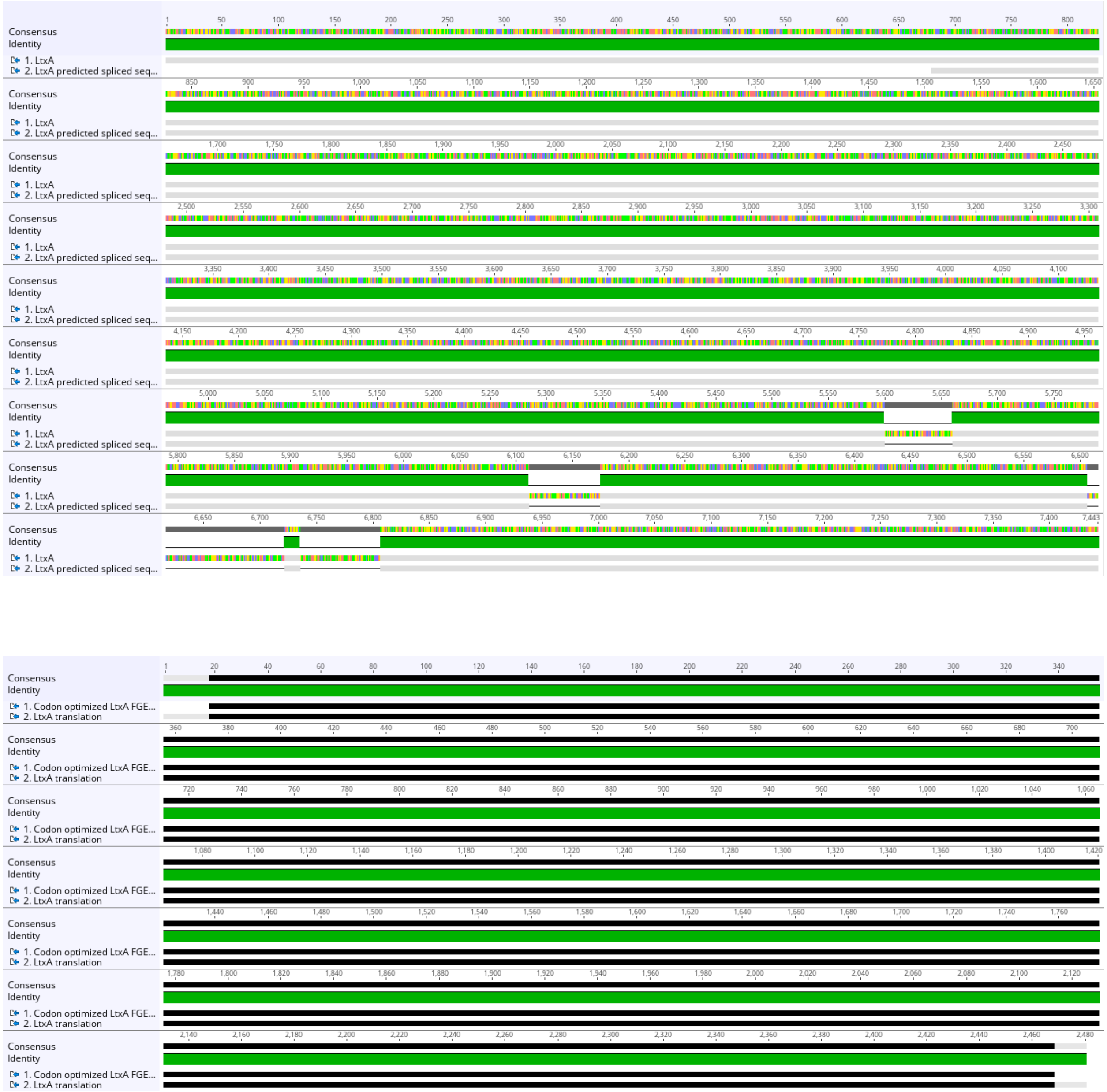

**Figure S9**: Sequence alignment of native *ltxA* sequence compared with FGENESH predicted sequence in *Aspergillus* sp. (top) showing predicted splice sites; sequence alignment of LTXA and codon-optimized LTXA predicted sequence from FGENESH (bottom) showing truncated sequence.

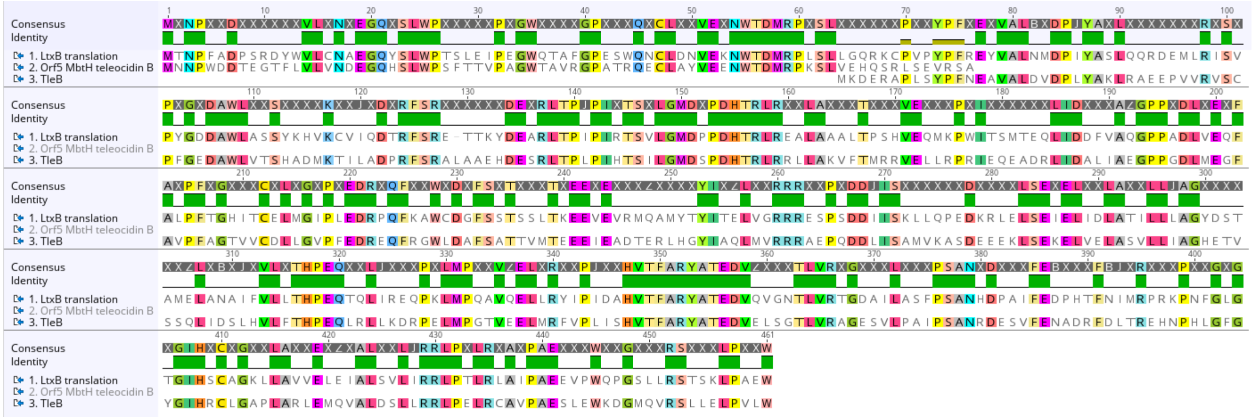

**Figure S10**: Sequence alignment of LtxB, TleB, and Orf5 (MbtH).

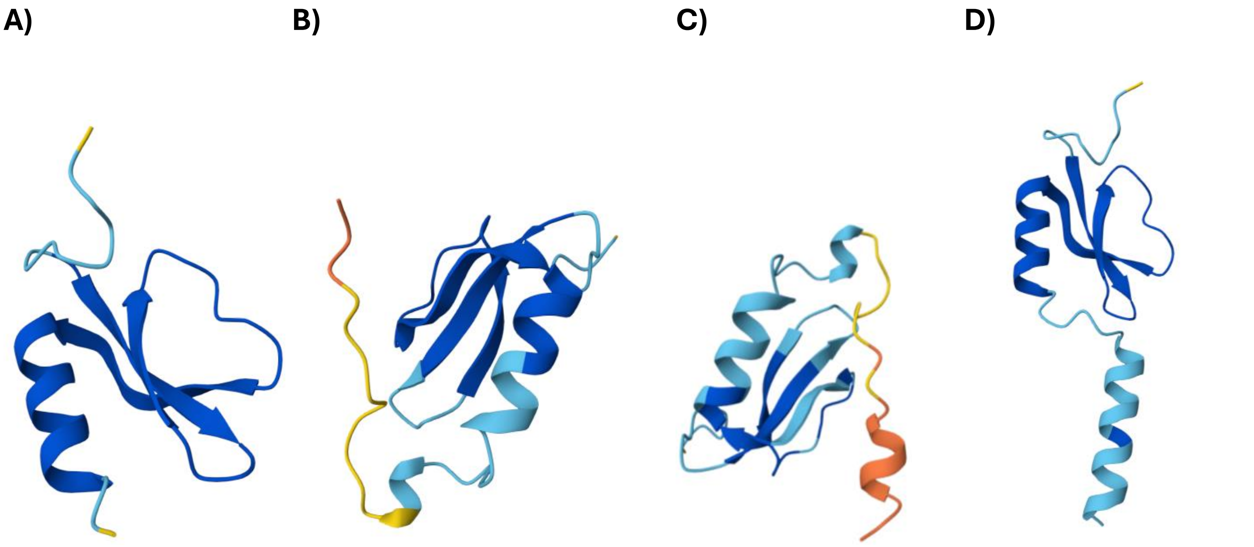

**Figure S11**: Models of the MbtH domain from LtxB using different sequence lengths (A-C) and comparison to Orf5 from the tle BGC (D).

#### 1.7 LCMS Methods

A HPLC-PDA-MS system was used for analysis of fungal extracts (Shimadzu LC2030C 3D Plus Prominence coupled to a Shimadzu LCMS-2020 mass spectrometer). Analyses were performed using a Phenomenex Kinetex $RP_{18}$ column (100 mm × 4.6 mm i.d., 2.6 $\mu$m) along with the Security Guard $RP_{18}$ protective guard column (4.6 mm i.d.) and eluting with $H_2O$ + 0.05% formic acid and MeCN + 0.05% formic acid using a gradient from 90:10 to 90:10 of $H_2O$-MeCN over 15 min, maintaining in 10:90 $H_2O$-MeCN for 3 min, from 10:90 to 90:10 in 1 min, and maintaining at 90:10 for 1 min, using a flow rate of 0.8 mL/min. The PDA detector scanned between $\lambda$ 190 and 700 nm. The MS was optimized using the following conditions: interface voltage 4.5 kV; interface temperature 350 °C; DL temperature 250 °C; heat block 200 °C; ESI mode, acquisition range 100 to 1000 Da; nebulizing gas 1.5 L $min^{-1}$; drying gas flow 15 L $min^{-1}$. Sample injection volumes was varied according to the analysis ranging from 1uL to 2.5 uL. were varied according to the analysis, ranging from 1 μL

**Direct Infusion HRMS/MS Analysis**

We first characterized selected metabolites using direct infusion high-resolution tandem mass spectrometry (HRMS/MS) on a SCIEX TripleTOF 6600+ mass spectrometer (AB SCIEX, Framingham, MA, USA). The sample was diluted 10-fold in 100% methanol (LCMS grade) and directly infused into the mass spectrometer at a flow rate of 10 μL/min to obtain high-quality MS/MS fragmentation spectra for structural annotation of the lyngbyatoxin standard.

Data were acquired in positive ion mode using Turbo Spray Ionization with an ion spray voltage of 5500 V. We set the declustering potential (DP), collision energy (CE), and collision energy spread (CES) to 80 V, 50 V, and 15 V, respectively. Nitrogen was used as curtain, nebulizing, and heater gas at 25 psi, 20 psi, and 15 psi, respectively. The source temperature was maintained at 25 °C. MS spectra were acquired with an accumulation time of 200 ms and processed using Analyst TF version 1.8.1 software (SCIEX).

**LC–MS/MS-Based Untargeted Analysis**

We subsequently performed LC–MS/MS-based untargeted analysis of WT and transformant extracts. Precursor (ILV) and final product (LTXA) standards were analyzed separately alongside the biological samples to support metabolite identification and validation. Analyses were carried out using an Exion ultra-high-performance liquid chromatography (UHPLC) system coupled to a SCIEX TripleTOF 6600+ high-resolution mass spectrometer (AB SCIEX, Framingham, MA, USA). The autosampler temperature was maintained at 15 °C throughout the analysis.

Metabolites were separated using a Phenomenex reversed-phase C18 analytical column (Kinetex 2.6 μm 100 Å C18, 100× 4.6 mm) coupled with Security Guard $RP_{18}$ protective guard column (4.6 mm). The mobile phases consisted of water containing 0.1% formic acid (Solvent A) and acetonitrile containing 0.1% formic acid (Solvent B). We used a flow rate of 0.8 mL/min with the following gradient: 0–0.5 min, 10% B; 0.5–15 min, 10–90% B; 15–18 min, 90% B; 18–18.1 min, 90–10% B; and 18.1–20 min, 10% B for column equilibration. The column temperature was maintained at 40 °C.

Untargeted LC–MS/MS data were acquired in positive ion mode using SWATH-MS data-independent acquisition (DIA). To generate variable SWATH windows, we first analyzed pooled quality control (QC) samples using data-dependent acquisition (DDA). MS survey scans were collected over an m/z range of 200–1000 with an accumulation time of 200 ms, while MS/MS spectra were acquired over an m/z range of 30–1000 with an accumulation time of 25 ms. The total cycle time was 1.15 s.

We set the electrospray ionization parameters as follows: ion spray voltage, 5000 V; curtain gas, 35 psi; nebulizer gas (GS1), 60 psi; heater gas (GS2), 80 psi; source temperature, 700 °C; declustering potential (DP), 50 V; collision energy (CE), 50 V; and collision energy spread (CES), 15 V. An APCI calibration solution was automatically infused every eight injections to correct for mass drift during the LC–MS/MS sequence. Data acquisition was performed using Analyst TF version 1.8.1 software (SCIEX).

#### Lyngbyatoxin calibration curve

The standard Lyngbyatoxin A was ordered from Cayman Chemicals with a purity of greater than 98%. The stock solution (0.5 mg/mL) was made. A dilution series was prepared to create a calibration curve for use in future PDA quantifications of Lyngbyatoxin A.

| Lyngbyatoxin A concentration (μg/mL) | Area under the curve |
|---|---|
| 12.5 | 505,282 |
| 6.25 | 245,929 |
| 3.12 | 118,767 |
| 1.56 | 59,255 |
| 0.78 | 27,686 |
| 0.39 | 13,002 |

**Table S9**: Serial dilutions of LTXA and corresponding peak areas used to create the calibration curve.

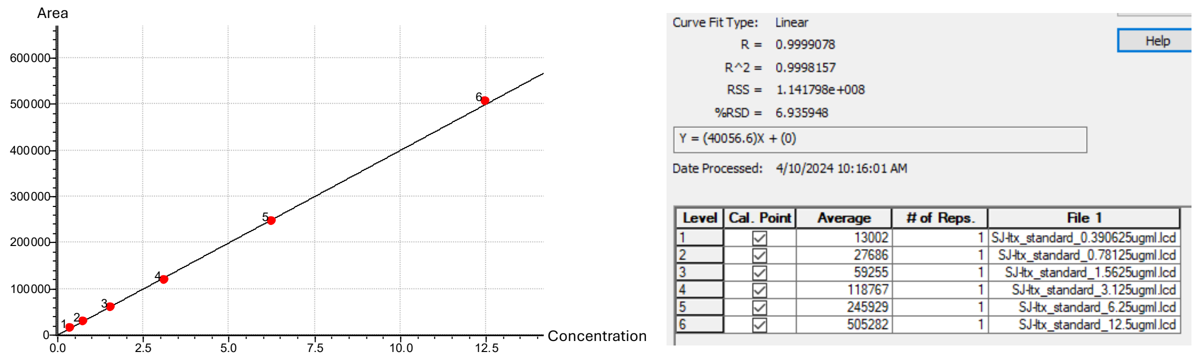

**Figure S12**: LTX calibration curve and linear fit curve calculated values.

#### 1.8 Protein expression, purification, analysis, and sequencing

*E. coli* BL21(DE3) cells transformed with the protein expression vector containing the targeted gene under the T7 promoter were streaked from glycerol stock onto LB agar media plates supplemented with 100 μg/mL ampicillin and incubated overnight at 37 °C. A single colony was used to inoculate 10 mL LB broth containing 100 µg/mL ampicillin, which was grown overnight at 37 °C with shaking at 220 rpm. The overnight culture was diluted 1:100 into 100 mL LB medium containing 100 µg/mL ampicillin in a 500 mL flask and grown at 37 °C with shaking at 220 rpm until the $OD_{600}$ reached 0.4–0.8. Protein expression was induced by adding isopropyl β-D-1-thiogalactopyranoside (IPTG) obtained from Invitrogen to a final concentration of 0.5 mM, and the culture was shifted to 25 °C for 18 hours with continued shaking. Cells were harvested by centrifugation at 4,000 × g for 30 minutes at 4 °C, and the pellet was stored at −80 °C until the extraction.

**6x-His tagged Protein Purification**

For purification, the *E. coli* cell pellet was resuspended in 10 mL of lysis buffer supplemented with a tablet of broad-spectrum protease inhibitor (Pierce Protease Inhibitor Mini Tablets, EDTA-free, Thermo Scientific). The cells were sonicated on ice (10 seconds on/10 seconds off, 10 minutes total on-time at 40% amplitude), and the lysate was clarified by centrifugation at 14,000 × g for 30 minutes at 4 °C. The supernatant was incubated with 2 mL Ni–NTA resin (equilibrated with buffer) at 4 °C for 1 hour with rotation, then loaded onto a gravity-flow column. The resin was washed with 100 mL of wash buffer-1, followed by 150 mL of wash buffer-2, and the His-tagged protein was eluted with elution buffer (20 mL).

**Protein Gel Electrophoresis (SDS-PAGE)**

SDS-PAGE gel was prepared using TGX™ FastCast™ Acrylamide Kit, 10% (Bio-Rad Laboratories, Hercules, CA). 5x Sample buffer (Boster's SDS PAGE Sample Buffer 5X (Reducing)) and 10x running buffer (Bio-Rad) were used according to the manufacturer's recommendations. ExcelBand Enhanced 3-color Regular Range Protein Marker (9–180 kDa, SMOBIO Technology Inc., Hsinchu City, Taiwan) was used as prestained protein ladder. The electrophoresis was run at a constant current of 140V for about 1 hour until the dye front reached the bottom of the gel. Then the gel is analyzed using Bio-Rad (GelDoc-XR).

Following electrophoresis, gels were rinsed in deionized water and equilibrated in transfer buffer for 15 minutes. PVDF membranes were activated in methanol for 1 minute, then equilibrated in transfer buffer for 5 minutes. The transfer sandwich was assembled by placing the equilibrated gel and the activated PVDF membrane between pre-soaked Western Blotting Filter Paper (Extra Thick, 8 cm x 13.5 cm) inside the transfer cassette of the Trans-Blot® Turbo™ Transfer System (Bio-Rad). The transfer program was selected from predefined mixed MW Midi (25V, 7 min).

Alternatively, the SDS-PAGE procedure was performed using hand-cast polyacrylamide gels with a 10 mL of 10% separating gel and a 4% stacking gel. Glass spacer plates were thoroughly cleaned, dried, and clamped together before use. For the 10% resolving gel, the 3.33mL of 30% acrylamide/bisacrylamide solution, 2.50 mL of 1.5 M Tris-HCl pH 8.8, 4.00 mL deionized water, 100uL of 10% SDS, 50 µL of freshly prepared 10% ammonium persulfate, and 15 µL of TEMED (N,N,N',N'-tetramethylethylene diamine) were combined in order gently to avoid introducing air bubbles, then immediately poured between the plates to the ~85% level of the glass plates. An IPA overlay was applied to create a flat interface and remove oxygen, and the gel was allowed to polymerize for 30 minutes. Once set, IPA was removed by pouring off, and the gel was dried by inserting a filter paper between 2 glass plates.

For the 5 mL 4% stacking gel, 2.00 mL of 30% acrylamide/bisacrylamide, 1.25 mL of 0.5 M Tris-HCl pH 6.8, 1.68 mL of deionized water, 50 µL of 10% SDS, 25 µL of 10% ammonium persulfate, and 15 µL of TEMED were mixed in order and poured above the resolving gel. A comb was inserted, and the stacking gel was left to polymerize for 30 minutes.

Protein samples were prepared by mixing with 5x sample buffer (Boster's SDS PAGE Sample Buffer 5X (Reducing)), then boiling for 5 minutes to ensure denaturation. After removing the comb, wells were rinsed with running buffer, and the gel was positioned in the electrophoresis tank. 1× running buffer (Bio-Rad) was used to fill the chambers. Samples and prestained protein markers (ExcelBand Enhanced 3-color Regular Range Protein Marker, 9-180 kDa, SMOBIO Technology Inc., Hsinchu City, Taiwan) were loaded, and electrophoresis was run at a constant current of 140V for about 1 hour until the dye front reached the bottom of the gel. Following electrophoresis, the gel was stained with Coomassie Blue stain by gently shaking it with the stain. Subsequently, the gel was destained with destaining solution by gentle shaking.

**Immunoblotting**

Following transfer, membranes were washed with 1× TBS for 5 minutes and blocked in 5% nonfat dry milk in TBS/T for 1 hour at room temperature with gentle agitation. Membranes were then washed three times for 5 minutes each with TBS/T. Primary antibody incubation was performed using 20 mL of His-tag antibody (6X His Epitope Tag Antibody, Rabbit Polyclonal, Rockland Immunochemicals Inc., Limerick, PA, USA) at a 1:2000 dilution in 10% blocking buffer (90% TBS/T) for overnight incubation at 4°C with gentle shaking. After three 5-minute washes with TBS/T, membranes were incubated with 20 mL of HRP-conjugated secondary antibody (diluted in 10% blocking buffer/90% TBS/T to get a 1:2000 dilution) for 1 hour at room temperature. Membranes were washed twice with TBS/T and once with 1× TBS for 5 minutes each

**Detection**

Protein bands were visualized using Clarity Western ECL Substrate (Bio-Rad) following 5 minutes of incubation at room temperature. Chemiluminescent signals were captured using a chemiluminescence imager, a colorimetric imager on Gel Doc XR+ Gel Documentation System, and two images were merged using Bio-Rad Image Lab software.

**Peptide Sequencing**

Protein bands detected in the de-stained SDS-PAGE gel were cut with a sterile blade, stored in ~100uL of 5% acetic acid, and shipped at room temperature, following the instructions provided by the MSU Proteomics facility. The peptide sequences obtained by MS-based sequencing were aligned with LtxB using Scaffold proteomics software.

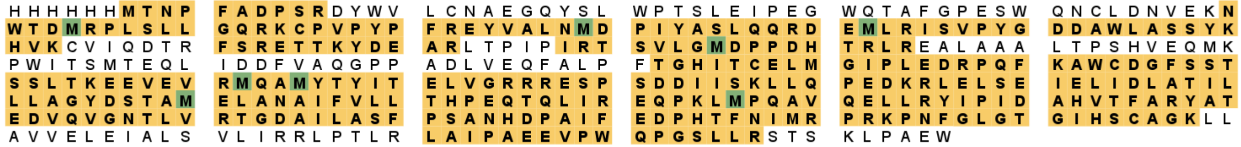

**Figure S13**: Sequencing results of N-terminal 6xHis-tagged LtxB.

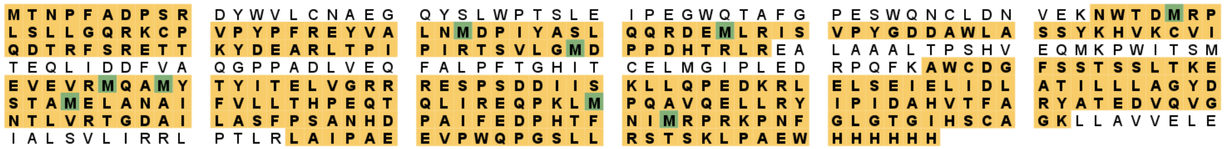

**Figure S14**: Sequencing results of C-terminal 6xHis-tagged LtxB.

### 2. LCMS traces

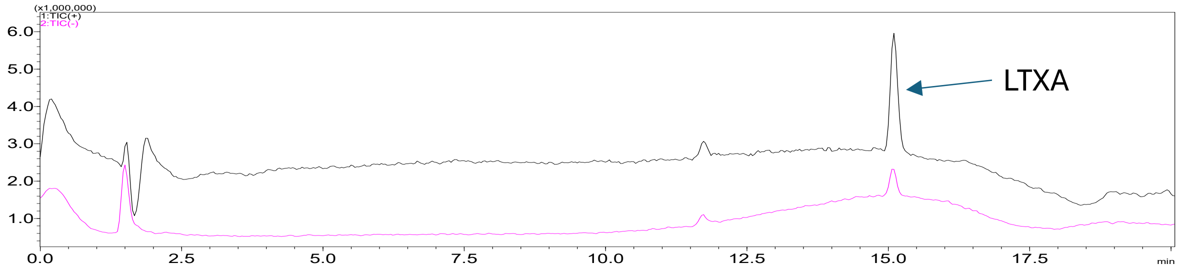

**Figure S15**: LCMS chromatogram of LTXA commercial standard.

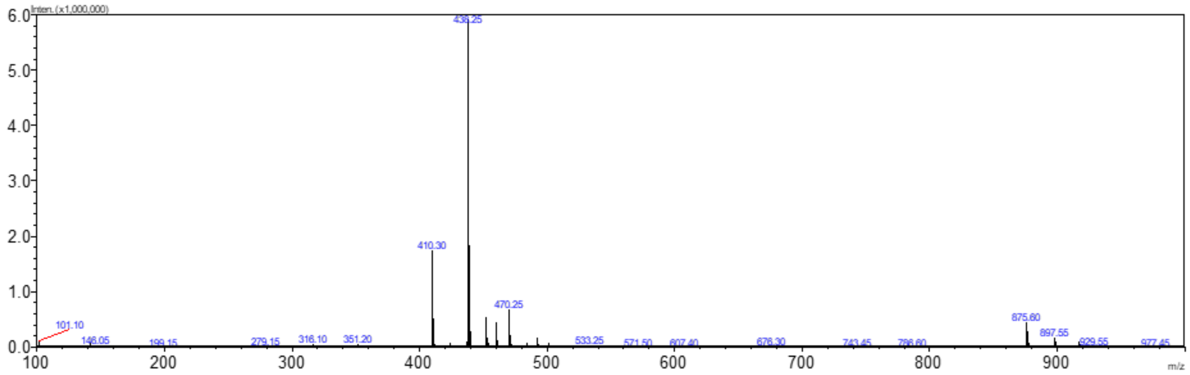

**Figure S16**: Low-resolution mass spectrum of LTXA commercial standard.

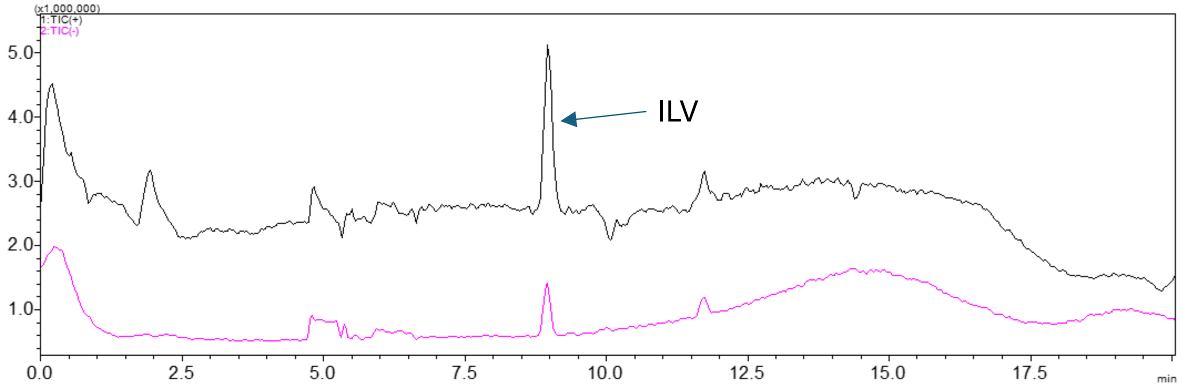

**Figure S17**: LCMS chromatogram of ILV commercial standard.

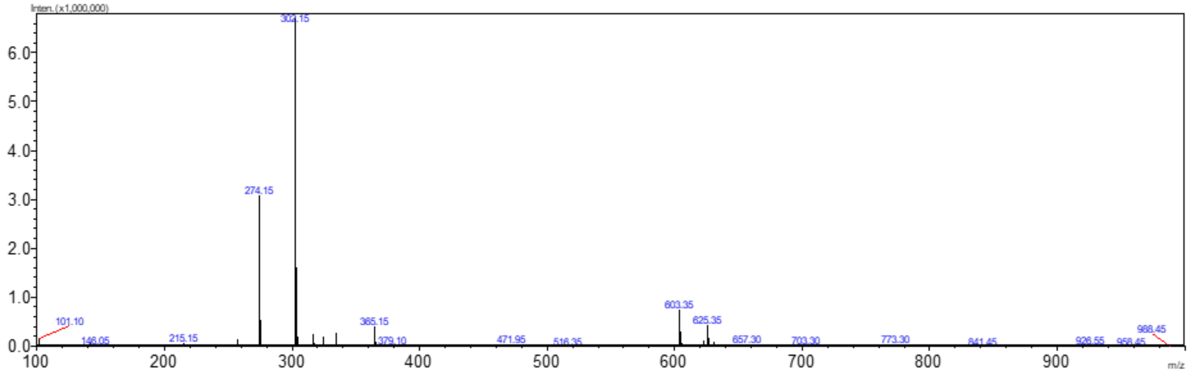

**Figure S18**: Low-resolution mass spectrum of ILV commercial standard.

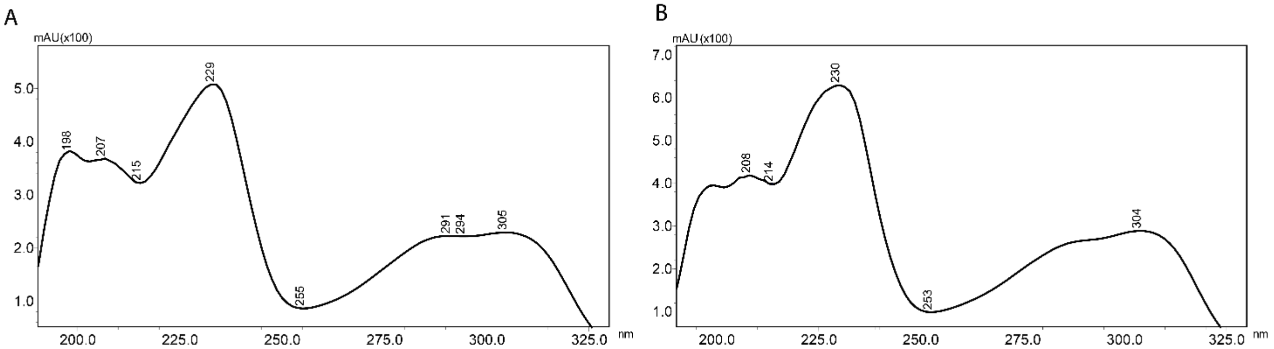

**Figure S19**: Absorption spectra of A) LTXA and B) ILV.

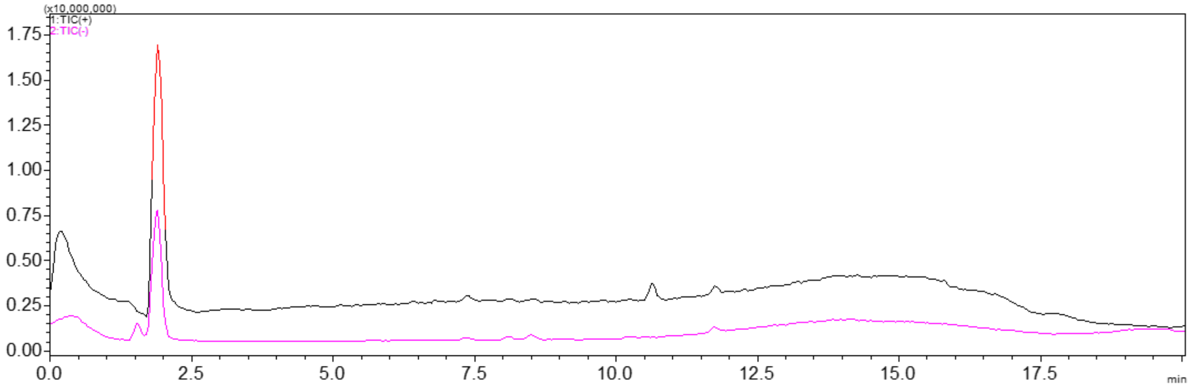

**Figure S20:** LCMS chromatogram of Val-Trp dipeptide commercial standard.

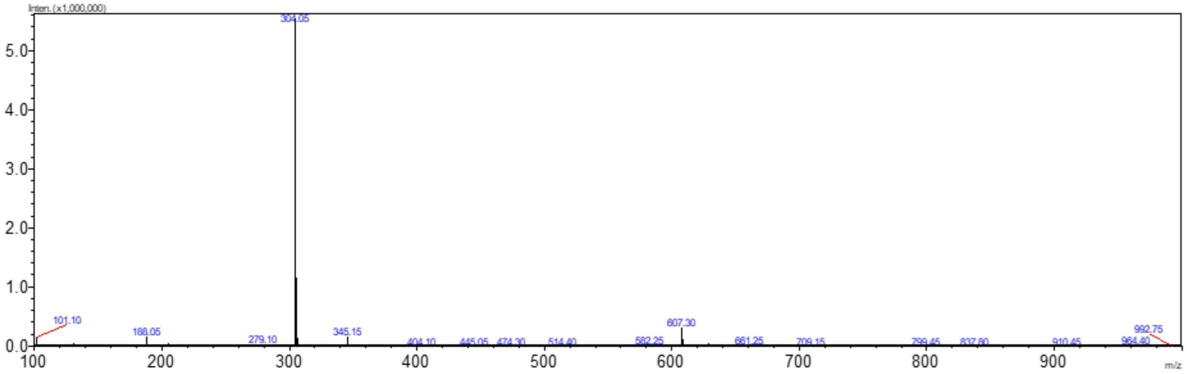

**Figure S21:** Low-resolution mass spectrum of Val-Trp dipeptide commercial standard.

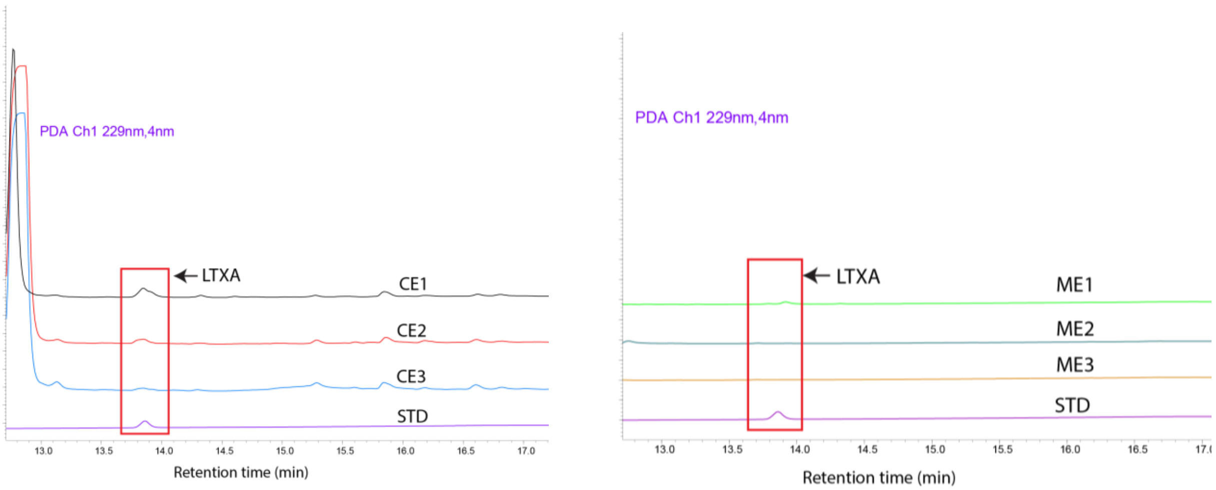

**Figure S22**: LCMS traces of *A. oryzae* NSAR1 cell extracts (CE) and media extracts (ME) after feeding with 25 µg of LTXA. A standard (STD) of LTXA at 12.5 µg/mL is included for reference.

| Sample | Calculated concentration of LTXA (µg/ mL) | Extraction efficiency (Conc (µg/ mL)*0.9 mL /25 µg)*100 |
|---|---|---|
| Growth media sample - 1 (ME1) | 3.55 | 12.78% |
| Growth media sample - 2 (ME2) | <0.3 | - |
| Growth media sample - 3 (ME3) | <0.3 | - |
| Cell extract sample - 1 (CE1) | 24.26 | 87.3% |
| Cell extract sample - 2 (CE2) | 26.58 | 95.6% |
| Cell extract sample - 3 (CE3) | 24.75 | 89.1% |

**Table S10**: Quantification and extraction efficiency of LTXA recovered from feeding experiments using the calibration curve from Figure S12.

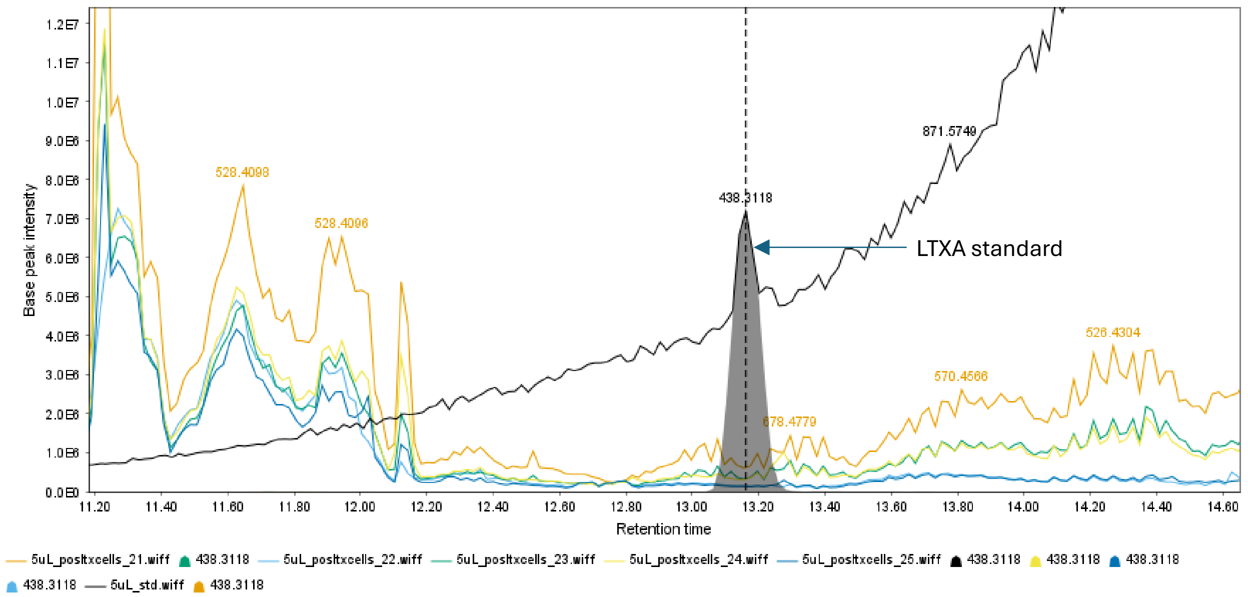

**Figure S23**: HR-LCMS analysis of *A. oryzae* + *ltxABC* + *hetI* cell extracts cultivated in various media (DPY = orange; CZD = green; CZD+starch = yellow; glucose = light blue; glucose-> starch = dark blue), compared to the LTXA standard ([M+H]+ *m/z* = 438.3118) shown in black.

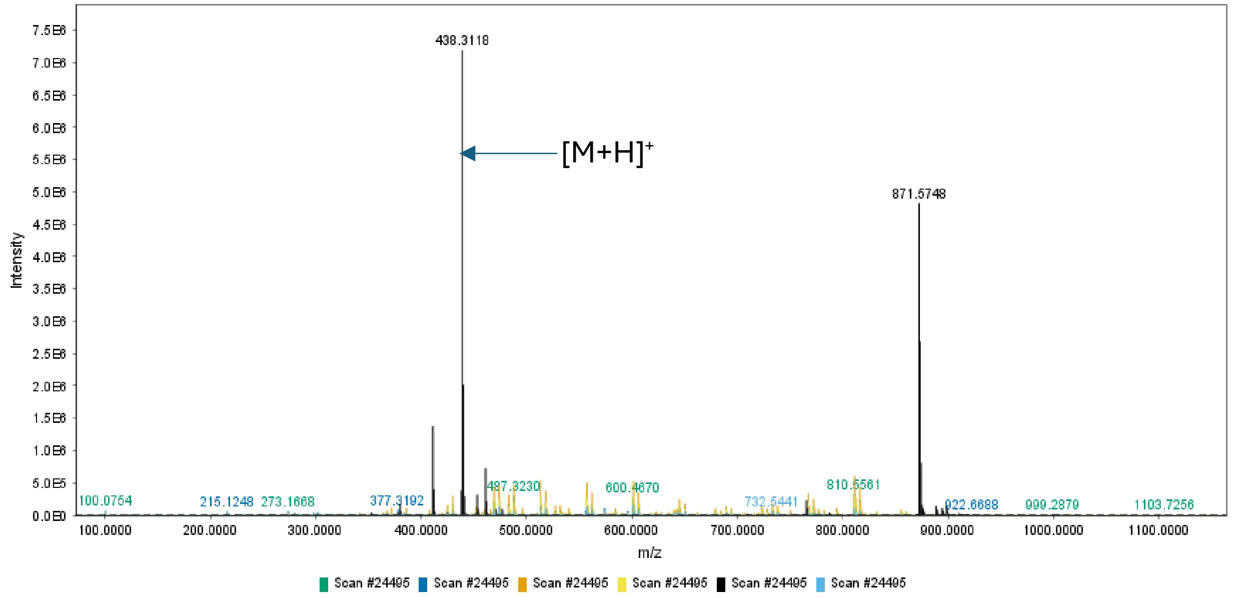

**Figure S24**: HRMS analysis of *A. oryzae* + *ltxABC* + *hetI* cell extracts cultivated in various media (DPY = orange; CZD = green; CZD+starch = yellow; glucose = light blue; glucose-> starch = dark blue), compared to the LTXA standard ([M+H]+ *m/z* = 438.3118) shown in black.

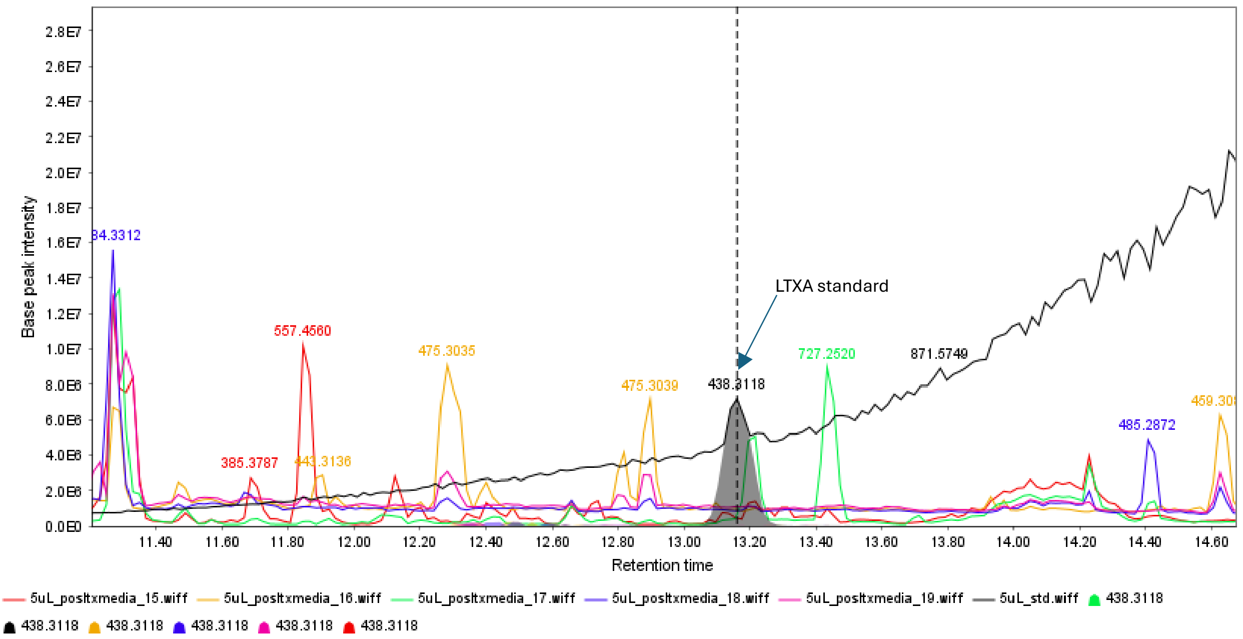

**Figure S25**: HR-LCMS analysis of *A. oryzae* + *ltxABC* + *hetI* media extracts cultivated in various media (DPY = red; CZD = green; CZD+starch = purple; glucose = orange; glucose-> starch = indigo), compared to the LTXA standard ([M+H]+ *m/z* = 438.3118) shown in black.

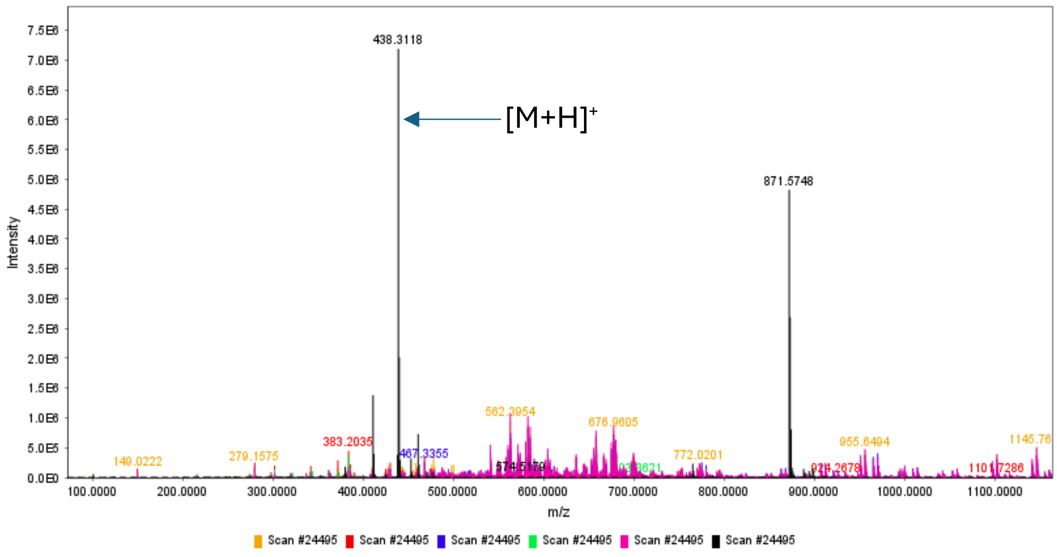

**Figure S26**: HRMS analysis of *A. oryzae* + *ltxABC* + *hetI* media extracts cultivated in various media (DPY = red; CZD = green; CZD+starch = purple; glucose = orange; glucose-> starch = indigo), compared to the LTXA standard ([M+H]+ *m/z* = 438.3118) shown in black.

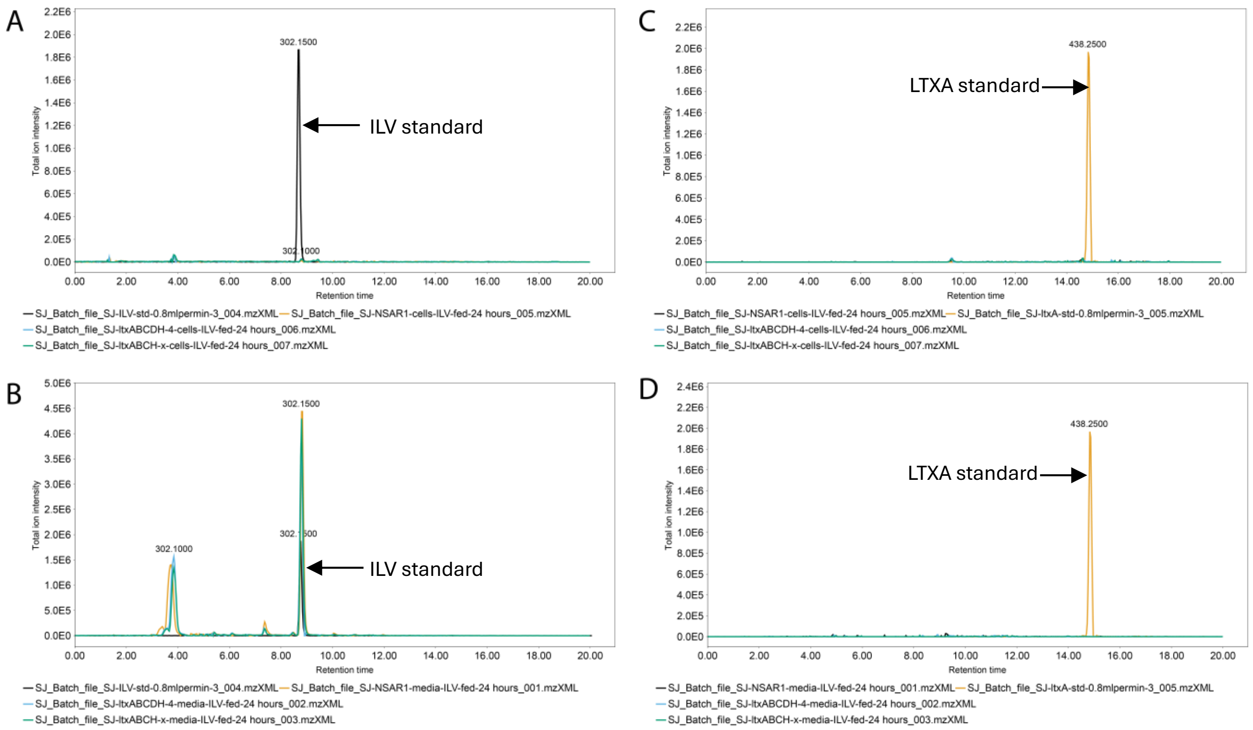

**Figure S27**: XICs of ILV and LTXA in the media and cell extracts of ILV fed to *A. oryzae* transformants 18,21. A) Cell extracts fed with ILV standard (black) showing negligible accumulation; B) media extracts fed with ILV standard (black) showing accumulation; C) Cell extracts fed with ILV showing no production of LTXA compared to a standard (yellow); D) media extracts fed with ILV showing no production of LTXA compared to a standard (yellow).

### References

#### Detecting Sequence Signals in Targeting Peptides Using Deep Learning

Abraham J, et al., Accurate structure prediction of biomolecular interactions with AlphaFold 3 (2024) 630: 493-500. https://doi.org/10.1038/s41586-024-07487-w

Armenteros JJA, Salvatore M, Winther O, Emanuelsson O, von Heijne G, Elofsson A, Nielsen H (2019) *Life Science Alliance* 2: e201900429. doi:10.26508/lsa.201900429

Horton P, Park KJ, Obayashi T, Fujita N, Harada H, Adams CJ, Nakai K (2007) WoLF PSORT: protein localization predictor. *Nucleic Acids Res*. 21: W585-W587. doi: 10.1093/nar/gkm259

Lazarus CM, Williams K, Bailey AM (2014) Reconstructing fungal natural product biosynthetic pathways. *Natural Product Reports*, 31:1339–1347. https://doi.org/10.1039/c4np00084f

Nakai K, Horton P (1999) PSORT: a program for detecting the sorting signals of proteins and predicting their subcellular localization, *Trends Biochem. Sci*, 24: 34-35. DOI: 10.1016/s0968-0004(98)01336-x

Pahirulzaman KAK, Williams K, Lazarus CM (2012) A toolkit for heterologous expression of metabolic pathways in aspergillus oryzae. In *Methods in Enzymology* 517: 241–260. Academic Press Inc. https://doi.org/10.1016/B978-0-12-404634-4.00012-7.
